# Frequent errors are the worst: robustness to individual failures in collective foraging

**DOI:** 10.64898/2026.07.12.738017

**Authors:** Chinmay Hemant Joshi, Anna Dornhaus

## Abstract

The collective behavior of complex systems emerges from the actions and interactions of their individual components. But what if these individuals make mistakes, or deviate from behavior tuned to lead to collective success? Here, we explore the effects of individual errors on collective outcomes. We investigate in particular whether information flow among individuals exacerbates or mitigates such individual failures. We use an agent-based, spatially explicit model inspired by collective foraging in social insects. Social insect colonies forage for food with autonomous workers who search for and exploit resources around the central nest, as well as share information about discovered resources. We find that the errors that have the most chances of occurring had the strongest impacts: for example, false positive detections can occur at any time during search, and each such error derailed exploration activity. Similarly, forgetting errors are potentially frequent and detrimental to resource exploitation. Despite the fact that communication inherently may narrow the breadth of information used by a colony, we found, in contrast, that it enhanced spatial exploration in our model. Communication in our model also reduced the effects of individual errors, instead of permitting erroneous information to spread. Our model thus illustrates that communication plays a central role in error management in complex systems, and that the evolution of communication systems in social insects may be shaped by selection on exploration and error mitigation as well as on efficient food retrieval.

## Introduction

In many complex adaptive systems, the behavior of individual units can only be understood in the context of their effect on the system as a whole. This is true, for example, for genes in a genome, cells in a multicellular organism, or workers in a social insect colony: in all cases, selection is primarily at the level of the collective [1–3]. Social insect colonies consist of a few reproductive queens and several (sometimes thousands) sterile workers that cooperatively perform various tasks such as foraging [4,5]. Even though queens control the reproduction in the colony, they do not have control over individual worker behavior; workers make independent decisions about where, when, and what to do, based on personal and social information [4–6]. In resource collection in particular, workers called foragers search for resources [7–10], and transport them to the nest [8]. They may communicate about the location, type, or quality of these resources with nestmates, and recruit others [6,11,12].

Like any other cognitive system, workers may make mistakes. These mistakes can stem from limitations in discerning rewarding and unrewarding stimuli [13], interference due to conflicting memories [14] or erroneous merging of related memories [15], inability to avoid responding to unrewarding stimuli due to innate biases [16], or noise causing errors in decision making [17]. We categorize the resulting potential errors based on the type of information affected: individual workers may falsely detect (or identify/classify) a non-existing resource (‘false positive’ [18]), or they may fail to detect an existing one (‘false negative’ [19]). Workers may forget an already-discovered resource location [20,21], or misremember it (thus returning to an incorrect location [14]). They may make an error in the sending or receiving of a communication signal (‘transmission error’[22,23]).

These errors can potentially extend well beyond the individual worker through communication, if erroneous information is spread through the colony. Such spread, or misinformation cascades, can lead to bad collective decisions [24,25]. Misinformation cascades and their effects are well documented across different animals, including humans [26–28]. Since communication in social insects can lead to positive feedback loops (amplification of and focus on a limited subset of information (Lanan et al., 2012)), they may be less likely to escape such bad decisions once a sizable proportion of the group is committed [30,25]. Erroneous information spread via communication signals may be particularly problematic if individual workers have fixed responses to stereotyped signals [31], and hence may have limited cognitive flexibility to ignore erroneous or outdated recruitment signals ([16]; but see: [32,33]. Communication and errors may alter that process in far-reaching ways, for example by symmetry breaking [34–36], altered exploration vs exploitation patterns [37–40], and uneven distribution of search effort [41,42].

However, social insects have also been shown to avoid or mitigate errors through a variety of ways. Workers can be flexible about both reacting to conspecific signals [43,44] and evaluating their reliability [45–47]. Workers may also repeatedly sample the same signal to reduce transmission error [48], or combine information from several sources (both personal [49,50], and social [51,52]). In addition, insect foragers tend not to amplify information unless they have individually verified it [43,44,53]. They may stop recruitment to suboptimal or dangerous locations by producing negative (repelling) signals [54,55], and by recruiting to alternative resource locations [53,56]. Therefore, the presence of communication among nestmates brings with it both the possibility of improved information collection and that of amplification of misinformation.

This paper uses a modeling approach to advance our understanding of how individual-level errors may affect collective performance. We investigated the effects of five different types of individual-level failures (false positive or false negative detection, wrong memory, forgetting, and transmission errors) on collective foraging. We employed an agent-based, spatially-explicit simulation model implemented in NetLogo to answer the following questions: (1) What effects do different types of errors have on different aspects of collective foraging performance? (2) Does communication mitigate or amplify the effects of individual errors?

To quantify the impact of individual errors on collective performance, one has to identify which dimensions of collective behavior matter. Here, we assume that the main objective of a foraging social insect colony is to maximize resource collection (i.e., rate of collection).

However, foraging consists of two qualitatively different activities: search and retrieval, often termed ‘exploration’ and ‘exploitation’ [9] . Exploration entails searching for resources and assessing their quality; exploitation implies collecting resources from known locations options (it is exploitation that is thought to primarily benefit from communication, although see Discussion). These two aims, search and retrieval, are typically seen as trading off with each other, as individuals can spend their time on only one of them [7,57], and colonies can allocate workers to one or the other activity [7,58,8]. The optimal allocation of time to exploration vs exploitation is the subject of much research [57,59,60] and is thought to depend, among other things, on the spatial and temporal distribution of resources in the environment [12,34,61]. To capture these different aspects of foraging, we separately quantify colony performance in exploration and exploitation as well as several metrics that contribute to these outcomes.

In the study of robustness to failures, what is often measured is not only performance at some arbitrary level of perturbation (or error frequency), but the ‘shape’ (functional form) of how performance declines with increasing perturbation or failure rate [62–65]. For example, a concave, initially slowly declining shape is sometimes called ‘graceful failure’ in engineering - one would want small problems to have small impact on overall performance, and only get a full system failure at high rate of component failure (whereas a convex shape implies failing components have an immediate, strong impact on system performance). We ask to what degree our simulated colonies achieve resilience up to high rates of different types of errors.

Social insect colony foraging has been previously modeled using individual-based simulations. This method is particularly suitable when exploring systems that depend on heterogeneity in space, time, or across individuals; all three of these are expected to be critical to social insect collective behavior. For example, individual-based models are frequently used to determine effects of various spatio-temporal resource distributions, particularly on benefits of communication [66–69,61,70]. Other models focus on spatial patterns in forager distributions, particularly in the context of symmetry breaking [29,71] or polydomy (the colony uses multiple nest sites) [72,41]. These and other individual-based models [73,74] have in common the different information states of individual foragers, the consequences of which are difficult to model with population-based models, and which also motivates our use of an individual-based model here.

Our aim here is to improve general understanding of the impact of different types of individual-level errors on complex system behavior; however, we used parameter values approximating empirical data from foraging social insect colonies, to capture the ecological and physiological constraints in this system.

## Results

### Impact of different error types on overall performance

The median number of resource units collected was lowered by all types of errors under all communication systems, but the magnitude of the performance decline differed among error types (Fig. 1, top six panels; p<0.001 in Kruskal-Wallis tests for each communication type, sample size range = 27929-50276 per test, Table S2). Pairwise comparisons in Dunn’s post hoc tests showed this to be a significant reduction in all but transmission errors, which caused the smallest median decline in performance, a decline which was only significant for the ‘beacon’ communication system. The strongest decline in resources collected was seen with false positive errors, which caused a median of 100% decline relative to the respective simulations with no errors (for all communication types); forgetting errors were similarly detrimental (median 83-97% decline), and false negative sensing and wrong target memory had intermediate effects.

**Fig. 1:**
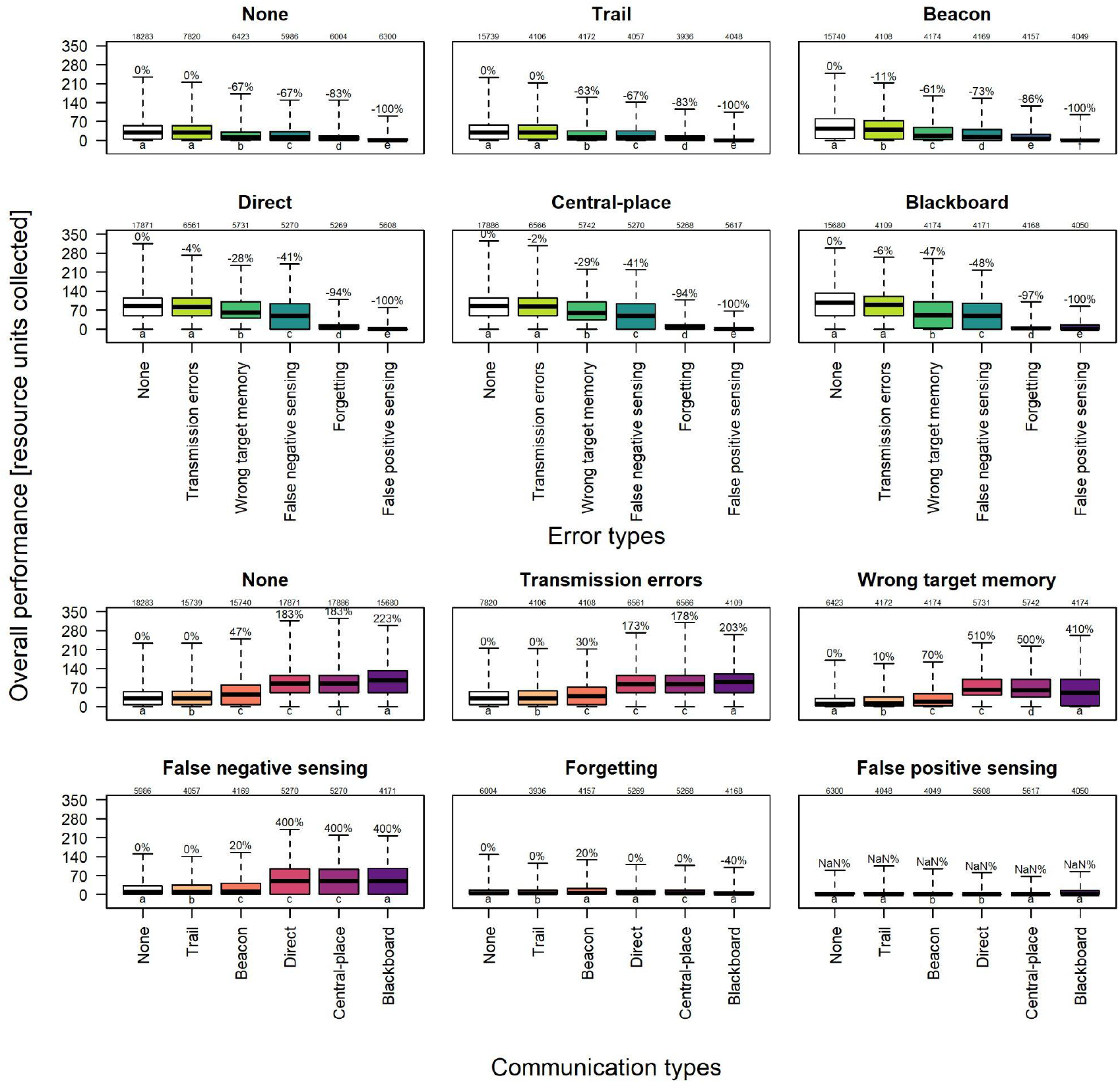
Total resources collected by the colony, separated by communication type and shown against error type in the top six panels; the bottom six panels show the same data, but separated by error type against communication type. Box plots show the median (solid black line), quartiles (colored box), and range (whiskers). Numbers on top of each panel indicate sample sizes (number of simulation runs). Letters at the bottom of each panel show Dunn’s post hoc test results (boxes that do not share the same letter are significantly different from each other, p<0.05 in pairwise comparison). Percentages above whiskers indicate relative change in median performance compared to median without error (top six panels) or without communication (bottom six panels) (white boxes). Errors generally led to decreased performance, while communication led to increased performance.

### The role of communication

When comparing these performance measures across communication systems instead of across error types, we find that for all error types, the communication type significantly affected the impact errors had on the number of resource units collected (Fig. 1, bottom six panels; p<0.001 for all Kruskal-Wallis tests, sample size range = 28802-99974 per test, Table S3). In most but not all cases, communication improved performance, sometimes dramatically (up to 510% compared to no communication). The near-total collapse of foraging with false positive sensing errors was not prevented by any communication type, and the impact of ‘trail’ communication was generally low (Dunn’s post hoc tests for individual pairwise comparisons). With forgetting errors, performance was also very low regardless of communication type, although here the only negative effect of communication was found (for the ‘blackboard’ type, where performance with communication was even worse than without). However, in all other cases, the effect of communication was positive, indicating that communication typically served to mitigate, not exacerbate, the impact of errors. With false negative sensing and wrong target memory errors, the benefit of communication was in fact even higher when errors were present than when they were not, and colonies with direct, central-place, or blackboard communication with these errors outperformed colonies with no errors and no communication. Overall, this shows that in this collective behavior scenario, misinformation cascades do not seem to impact performance, if they occur at all. Instead, communication rescues collective outcomes from individual errors.

### The shape of robustness

The impact of errors as reported above was evaluated for a single, standard magnitude and probability of errors. However, an important trait of robust systems is their ability to fail ‘gracefully’, i.e., prevent performance collapse at small error rates (instead ideally declining gradually and initially slowly with increasing failures) [62,65]. We thus evaluated the performance of colonies with increasing error probability, for all communication and error types (Fig. 2). All error types led to a reduction in resource units collected with increasing error probability (with the exception of transmission errors). However, the two most detrimental error types, false positive sensing and forgetting errors, show an immediate steep decay in performance (in the convex region of the S-curve over the entire error probability range), whereas wrong target memory and false negative errors show an initially slow decline, and transmission errors, if showing a decline at all, remain in the concave region of the S-curve. This pattern remained qualitatively the same across all communication types.

**Fig. 2:**
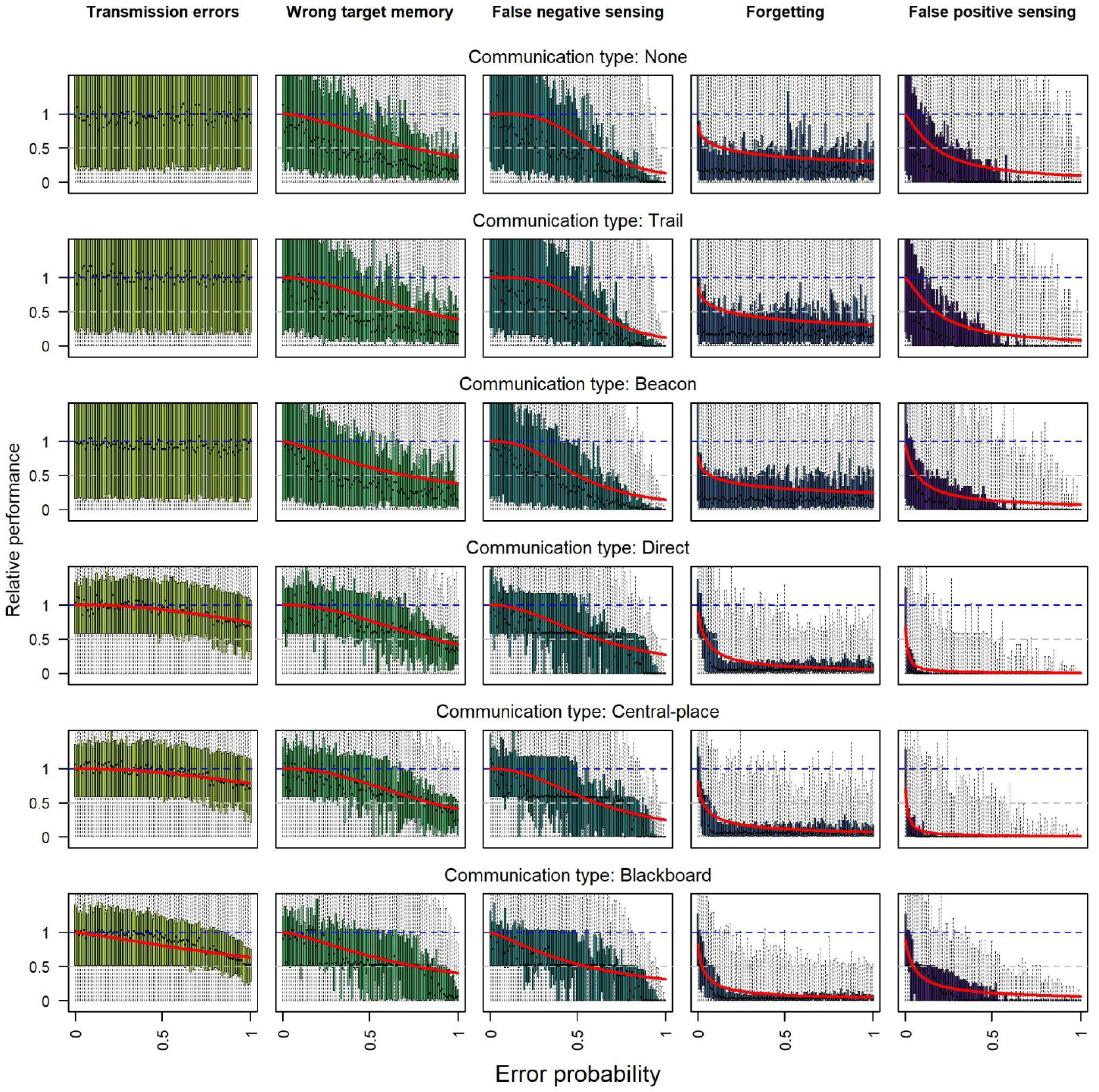
Relative performance (resource units collected relative to the median resource units collected with no errors) plotted against error probability for each combination of error and communication type (as boxplots with 100 bins from 0 to 1). Red lines indicate fits of inverted hill function (S curve starting at 1 and decreasing) to the raw data. The ‘shape’ of the fit indicates whether performance declined quickly even for minimal errors, or remained steady up to higher error frequencies. False positive and forgetting errors show a steep, convex drop in performance, while transmission errors show a minimal, concave decrease. Wrong target memory and false negative errors exhibit a complete S-curve. The estimation process did not converge for transmission errors for no communication, beacon and trail communication, presumably because performance either initially increased or did not drop over the range of error probabilities (by definition no drop should be present for transmission errors under no communication, top left panel).

### Exploration vs. exploitation

To better understand how errors impacted colony performance, we evaluated colonies both in terms of their ‘exploration’ and ‘exploitation’ (or search and retrieval) [9,57]. Exploration can be framed as a combination of investment in search (i.e., forager-time spent searching), area allocation or avoidance of overlap between searchers (i.e., amount of new area explored given a particular investment in search), and search efficiency (ability to actually discover resource patches given a particular area searched). These three measures combine to give the total number of resource patches discovered, a measure of successful search (Fig. 3). For all these performance measures, across all communication systems, error types differed in their impact (all Kruskal-Wallis tests p<0.05; Tables S4-S7; see Dunn’s post hoc tests for pairwise comparisons, Fig. 3). The severity order of the error types for exploration remains the same as for overall performance (compare Fig. 1 & Fig. 3, left column), even though some errors actually improved mutual avoidance of searchers (3rd column in Fig. 3) and investment in search (4th column in Fig. 3). Interestingly, despite the fact that overall performance is similarly impacted by false positive sensing and forgetting errors, false positive sensing clearly has the strongest impact on exploration: with this error type, investment in search drops to almost nothing (Fig. 3, right column, rightmost box), and so does search success (number of resources discovered, leftmost column).

**Fig. 3:**
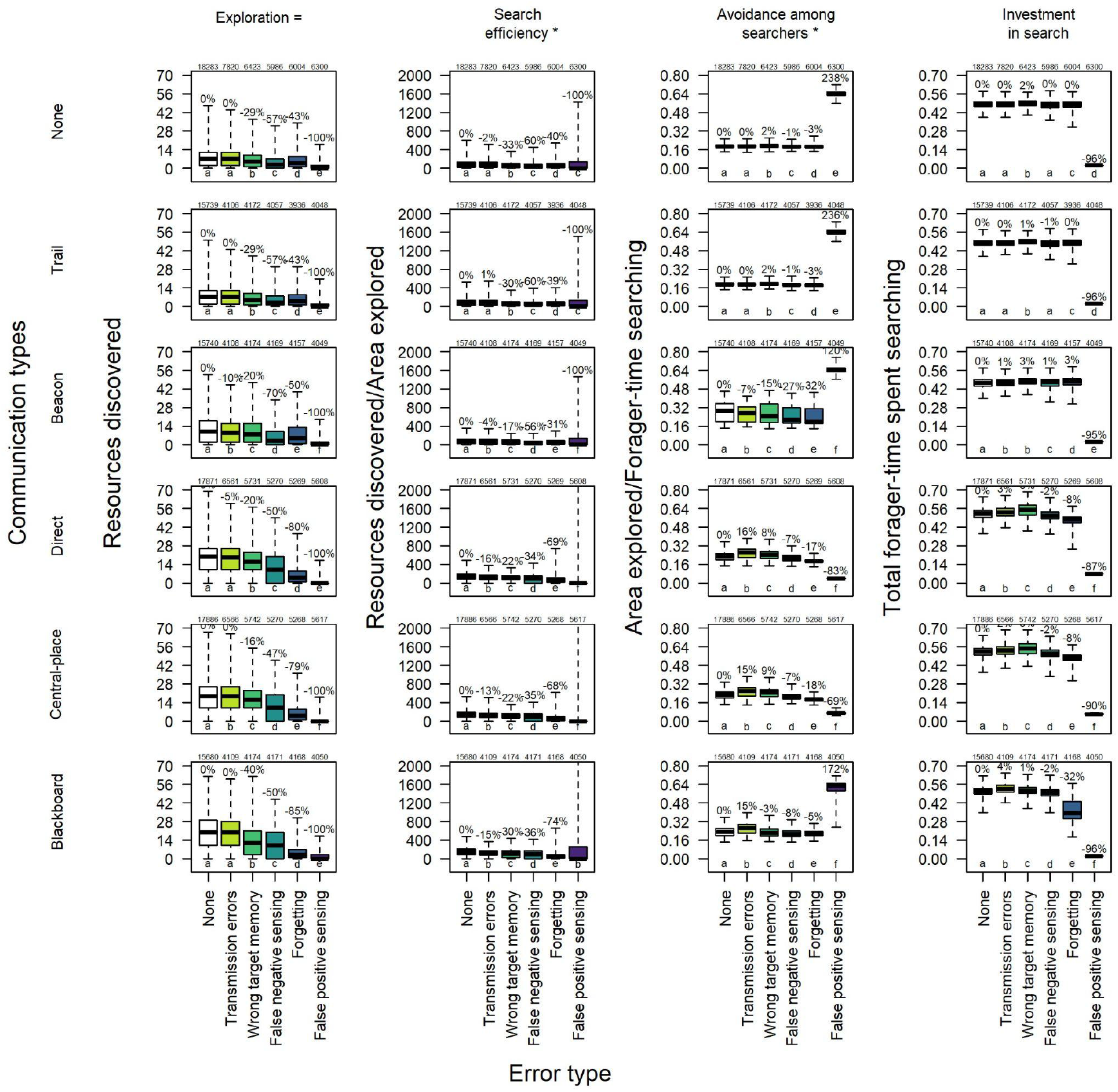
Ability of colonies to explore the environment, here measured as number of resource patches discovered (Exploration), which is a product of Search efficiency (resource patches discovered/proportion of total area explored), Avoidance among searchers (proportion of total area explored/proportion of time spent searching), and Investment in search (proportion of time spent searching). Box plots show the median (solid black line), quartiles (colored box), and range (whiskers). Numbers on top of each panel indicate sample sizes (number of simulation runs). Letters at the bottom of each panel show Dunn’s post hoc test results (boxes that do not share the same letter are significantly different from each other, p<0.05 in pairwise comparison). Percentages above whiskers indicate relative change in median performance compared to median without error (white boxes). The map contained a total of 100 resource patches. Errors severely negatively impact exploration, particularly in search efficiency; false positive sensing errors also strongly impact investment in search.

Exploitation (or retrieval, measured as the amount of resource units collected per resource patches discovered) is the product of investment in retrieval (i.e., forager-time spent collecting) and collection efficiency (i.e., amount of resource units retrieved per time invested) (Fig. 4). In our simulation, the time invested in retrieval was much lower than the time invested in search (compare the y-axis units in the right columns in Fig. 3 and 4; also see Fig. S1 and S2 in supplementary material). The overall exploitation performance across error types differed from the order in exploration and overall performance impacts (all Kruskal-Wallis tests p<0.05; Tables S8-S10; see Dunn’s post hoc tests for pairwise comparisons, Fig. 4). In particular, forgetting impacted exploitation the strongest, and wrong target memory also had a higher impact on exploitation, compared to other error types, than on exploration. In addition, communication types interacted with error types to affect their impact.

**Fig. 4:**
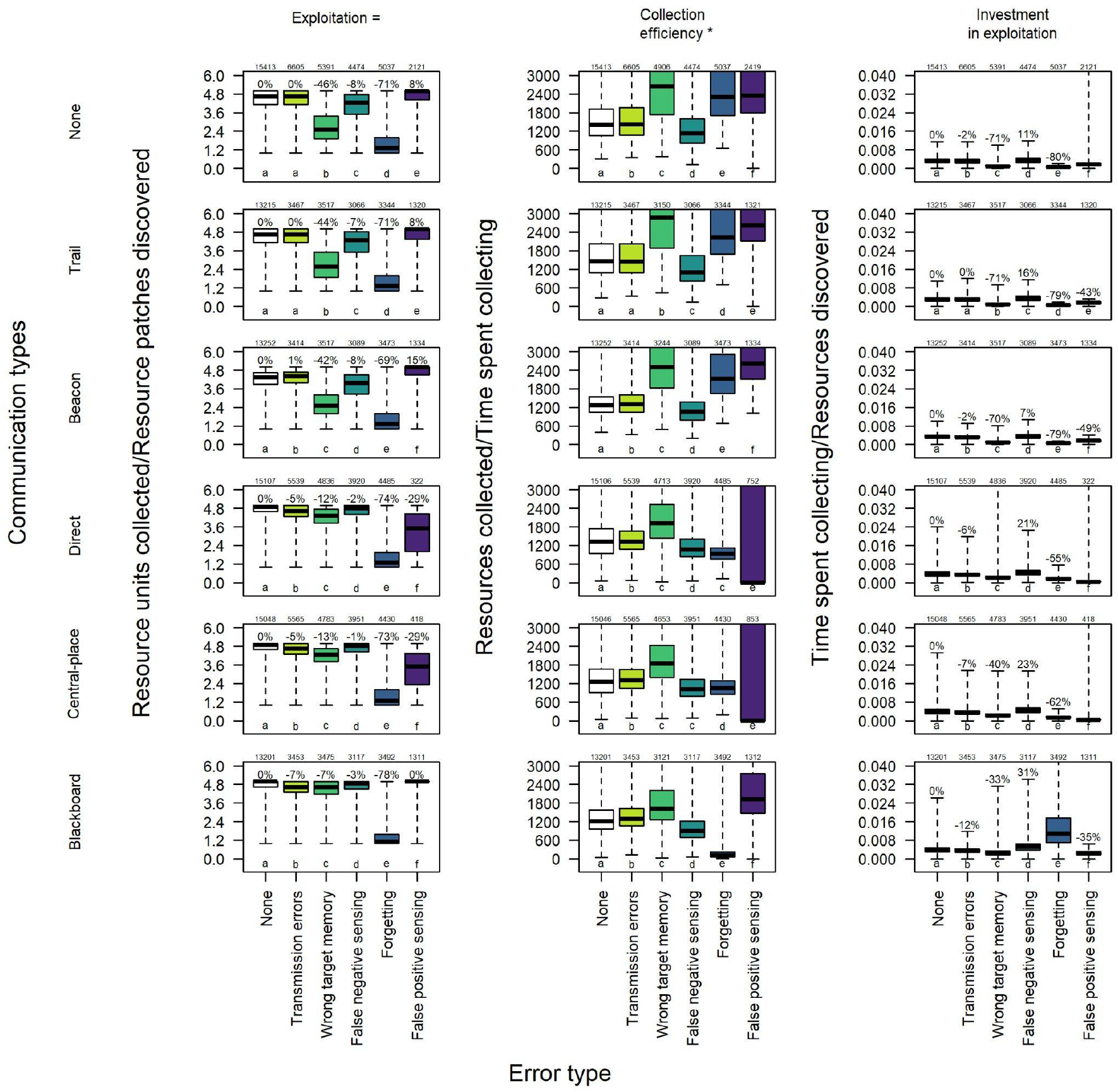
Ability of colonies to exploit discovered resources, here measured as collected resource units/resource patches discovered (Exploitation), which is a product of Collection efficiency (resource units collected/proportion of time spent collecting) and Investment in exploitation (proportion of time spent collecting). Box plots show the median (solid black line), quartiles (colored box), and range (whiskers). Numbers on top of each panel indicate sample sizes (number of simulation runs). Letters at the bottom of each panel show Dunn’s post hoc test results (boxes that do not share the same letter are significantly different from each other, p<0.05 in pairwise comparison). Percentages above whiskers indicate relative change in median performance compared to median without error (white boxes). Each resource patch contained 5 resource units (thus maximum collection efficiency is 5). The impact of different error types on exploitation is very different from that on exploration; forgetting most strongly impacts resource collection efficiency. The time investment in collection is much lower than that in search.

### What drives the impact of errors?

What mechanisms can explain these impacts? Overall, the number of resource units collected is tightly predicted by the number of resources discovered (Fig. 5, panel h), indicating that with most errors (and also most communication systems, Figs. S3-7), foraging performance is limited by exploration more than exploitation in our model. Some variation is also explained by the stochastic variation on how close resources are placed to the nest (panel d, note the overall negative slope) and how quickly the first resource is discovered (negative slope in panel b - this is a result of a combination of stochastic variation in search paths and resource distribution, and the search efficiency of colonies with particular error and communication types).

**Fig. 5:**
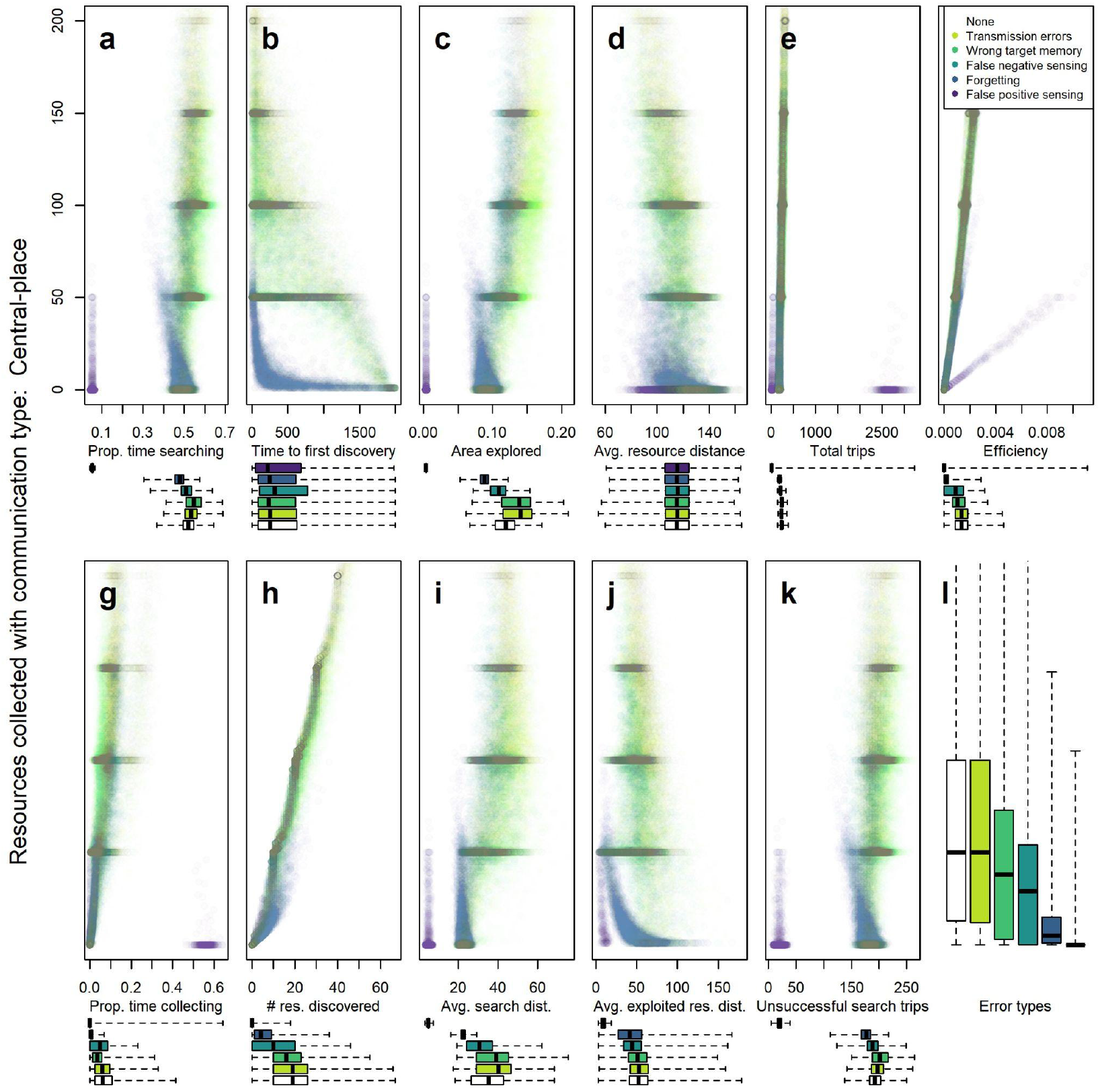
Example, for the central-place communication type, of the impact of errors on different measures of colony performance and behavior (for other communication types see supplementary materials). Boxplots underneath scatterplots show x-axis value distributions for each error type; y-axis (performance) distributions for each error type are shown in panel I (and are identical across all scatterplots; panel I matches corresponding data in Fig. 1(top six panels)). Dark horizontal lines show high point densities; since resources occurred in ‘clusters’ of 10 resource patches carrying 5 resource units each, colonies that emptied exactly one resource cluster collected 50 resource units (or multiples of 50 for multiple clusters). Intermediate values indicate simulation runs in which some resource clusters were not fully exploited.

The two most detrimental errors show distinct differences in collective behavior. While both false positive sensing and forgetting achieve much lower success (common y-axis in Fig. 5) even for a similar range of time to first discovery, false positive sensing has foragers spend a lot of time attempting to collect resources (panel g), despite almost no resources being discovered (panel h). The false positive sensing error implies that foragers who are searching may ‘detect’ a resource that is not there, and return to the nest with this ‘target’ in their memory. On returning to the location, they fail to find anything at the location, and resume exploring. This error thus leads to frequently short search trips (purple points in Fig. 5, panel a) and thus short search distances (panel i) and little area explored (panel c). There are many (attempted) collection trips (panel e), but little to show for it (since there was no resource there in the first place, panel g), and any of the few resources that are found (panel h) are a short distance from the nest (panel j); the same issues occur with or without communication (compare to Fig. S3). Forgetting errors on the other hand generate a similarly strong impact, but by a different mechanism: foragers who already have information about a resource ‘forget’ this information and enter the exploring state. This leads to a lower number of resource units collected per resource discovered (blue points in Fig. 5, panel h; also see ‘forgetting’ box in left column in Fig. 4). It is also notable that the range of time spent searching (panel a), area explored (panel c), and exploited resource distance (panel j) for forgetting is not dramatically different from errors that lead to much lower impact - however, with other errors, more time searching and searching farther away positively correlate with foraging success (but do not when foragers ‘forget’ about these discovered resources). In addition, colonies are less able to capitalize on communication (all outcomes for forgetting look more similar to the ones without communication: compare blue points in Fig. 5 to Fig. S3).

## Discussion

Individuals making errors are detrimental to collective success; but we found that the exact type of error that individuals make radically affects its impact. Not only is this true quantitatively (in that some error types are more detrimental), but also qualitatively: first, some errors cause more ‘graceful’, i.e., gradual, failure whereas others lead quickly to collapse in performance; and second, different errors have impacts in different parts of the process, affecting its interaction with the environment as well as with communication strategy. We also asked whether communication in general is likely to exacerbate or mitigate the effect of errors on collective performance, given its potential to amplify both correct information and misinformation. We found that communication almost always improved foraging success, including in the presence of errors. Moreover, in some cases, communication had an even bigger impact when errors were present, thus mitigating the negative impact of errors. Lastly, we found several surprising effects we had not specifically looked for. First, errors did not generally increase effective exploration, nor did communication reduce exploration. Instead, the relationship between exploration and exploitation was generally positive (Fig. S1). Second, in a perhaps related finding, foraging success was almost entirely explained by ability to discover resources, and the actual collection of food from ‘known’ resources was apparently neither costly nor difficult unless foragers made memory-related errors. Third, despite the strong positive effect of communication, there was barely an effect of transmission errors, indicating perhaps that the benefit of communication was driven more by its effect on activating foragers and drawing them further from the nest rather than in precise communication of location. We discuss each of these findings in detail below.

### Error types that have more chances of occurring have more impact

What explains the differences in impact across the different error types? We had used the same ‘error probability’ (per individual and timestep) for each error type. However, this does not mean that all errors occurred equally frequently overall, since each error could only occur when foragers were in a specific state. Foragers could falsely detect a food source (false positive errors) every timestep they were in the ‘exploring’ state; they could forget a remembered food location every time step in the ‘inactive’, ‘exploiting’, or ‘returning forager’ states. However, false negative detections could only occur when an ‘exploring’ forager actually came in sensing range of a resource patch, transmission errors only occurred on signaling, and ‘wrong target memory’ only occurred at the moment that foragers committed a new resource location to memory. Foragers spent significantly more time in the ‘exploring’ than the ‘exploiting’ (or any other active) state (Figs. 4 & 5, right columns), meaning that the actual number of times a ‘false positive detection’ occurred was likely much higher than that for any other error. Wrong target memory errors, on the other hand, were likely rarer than other errors. It is striking that the two most detrimental errors, false positive sensing and forgetting, are the ones that can occur in the most common behavioral states, ‘exploring’ and ‘inactive’. In our simulations, the impact of errors on the amount of resource collected was thus likely explained in large part by such differences in the number of times each error occurred.

Decisions that occur frequently are thus likely to be under stronger selection to get right than decisions that occur rarely, everything else being equal [88,89]. With respect to errors during foraging, false positive and forgetting errors are therefore more likely to be either avoided and/or mitigated in foraging animals than, for example, false negative sensing, wrong target memory, or transmission errors. There is an inherent tradeoff in minimizing false positive and false negative errors in any decision-making system [13]. Animals minimize one error type over another depending on the cost incurred due to these errors. For example, individuals in a group initiate false alarm signals at a higher rate to warn other group members when it is difficult to detect predators (under cloudy or low light conditions for instance, almost 4-6 false signals over 1 correct signal) [26,28], or avoid valuable food sources that typically have cryptic ambush predators (5% false positive rates in bees erroneously avoiding visiting flowers of a certain color/type that were associated with cryptic predators in the past, but do not have the predators at present [90,91]). Such false alarms can be costly in terms of costs of responding, reduced foraging efficiency, or opportunity cost [28,91]; but failing to react when a predator is present can lead to total fitness loss. In such scenarios, animals are thought to prioritize avoiding false negative errors over false positive ones [92]. However, our results are a reminder that for the same probability of a particular error occurring, given the right condition is present, if the item to be detected is actually rare, false positive errors are likely to vastly outnumber false negative ones.

While ‘frequent errors are the worst’ may seem somewhat obvious in hindsight, this is exactly the kind of insight that models, by forcing clearer, more quantitative thinking, can provide [93,94]. In fact it is based on the same phenomenon as the ‘medical test paradox’ (referring to the fact that false positives of a diagnostic test may vastly outnumber true positives when a disease is generally rare) and the claim that ‘most research findings are false’ [95]. In all these cases, the tradeoff between false positive and false negative errors is shaped by the base probability of actual positive or negative events.

### Robustness shape is characteristic of a given error type

Robustness analysis allows us to quantify the system’s performance against increasing levels of perturbations. We generally define performance to be at 100% without errors, and expect it to decline to some low level at maximal error rate (typically to 0%, unless the function affected by the errors is not essential for overall performance). However, the ‘shape’ of this decline, i.e., how early and steeply performance decreases with increasing component failure rates, characterizes the robustness of a system and ultimately its ability to withstand typical levels of perturbation [62–65]. By quantifying the shape of a social insect colony’s robustness to individual errors, we illustrate the impact of such errors beyond any specific assumed error probability (which we had arbitrarily, and highly, set to 0.5 for most of the presented results). We show that for the most detrimental errors, performance declines steeply even for very low error probabilities, whereas system performance stays flat or declines only very gradually up to high error probabilities for other types of error (Fig. 2).

### Communication mitigated the effect of errors

Information exchange is broadly seen as one of the primary benefits of high individual density [96], and one of the key adaptations that make social insect colonies successful [6,97,98]. Communication in foraging in social insects is particularly well studied [6,66,99,100]. However, the benefits of communication are surprisingly not always obvious [66,100,101]. Empirical [100,102,103], and modeling [104,105] studies have found a variety of costs of communication. A particularly general effect of communication is the potential for a narrowed spectrum of resources used (which may represent a benefit [103,106] or a cost [66,107]), or which results in symmetry breaking in extreme cases [29,36]. Another possible cost of communication is the amplification of incorrect information, or ‘misinformation cascades’ [25,108]. The conditions in which misinformation spreads through social networks have received much attention in recent literature (e.g., [109,110]. Empirical studies have shown that social insects use a variety of mechanisms to evaluate or improve the quality of social information, from evaluating the reliability of multiple information sources and strong sensitivity to unreliable signallers [45,47] to integrating over several signallers before important decisions with quorum thresholds [111]. Social insects generally also always personally evaluate information before further communicating it [48,44,53], and may use repellent signals to counteract what they perceive to be erroneous information [54,55].

However, in our study, we did not code any of these more sophisticated mechanisms for counteracting or buffering misinformation and misinformation cascades; we were interested in quantifying the potential impact of amplification of individual errors in the basic case of collective foraging with communication. Despite this, we did not find that communication decreased performance when individuals were error-prone, demonstrating that in this case, the mitigating effects of social information can be solely attributed to the fundamental design of the communication system itself.

### Errors do not universally increase exploration

Individual errors, in the foraging literature, are often considered to be synonymous with choosing exploration over exploitation [59,60]. Exploration could involve either choosing a resource presumed to be inferior (typically designated as ‘sampling’ [60]) or movement to a new or previously unrewarding location (typically designated as search - [57,60]). Either behavior is similar to ‘making errors’ in that the forager is not visiting the resource currently estimated to be most profitable. However, some degree of exploration is part of an optimal foraging strategy as it is necessary to collect (and update) information about the environment and the best options [9,57,60]. Similarly, errors in extracting information from communication signals are sometimes thought to be adaptive in preventing symmetry breaking and improving the breadth of resource use [29,112].

Here, however, we show that errors in many cases may in fact be universally detrimental to foraging, by causing time or energy costs that prevent foragers both from collecting food and collecting information. We measured several aspects of exploration (Fig. 3 and Figs 6 & S3-7). Almost universally, errors decreased performance even in terms of exploration, with one exception: transmission and wrong target memory errors (with blackboard, central-place, or direct communication) do seem to increase area explored per time invested, probably by sending searchers to new areas - perhaps similar to what had been proposed for honey bees in the ‘tuned error hypothesis’ [112]. Time invested in searching is also higher in several cases with errors compared to without errors. The reason that errors don’t generally increase exploration appears to be that most errors, instead of allowing foragers to explore novel areas, interrupt trips and cause foragers to stay closer to the nest, an area that tends to already have been thoroughly explored. In the case of wrong target memory, foragers instead are committed to reaching a particular patch, which may be some distance from the nest. Even when this specific patch does not contain a resource, this error thus draws foragers some distance from the nest, increasing exploration. The overall impact on foraging success is still negative in our simulations, presumably because resource patches already discovered are not fully exploited (panel h in Fig. 6 and S3-7), at least for the combination of error probability and resource distribution simulated here. As stated above and further discussed below, transmission errors had a very low impact across the board in our simulations. However, there is some indication that they also have the potential to increase exploration (e.g. increased search distance and area explored, panels c, h, and i in Fig. 6 and S3-7), as previously proposed in the literature [29,112].

**Fig. 6:**
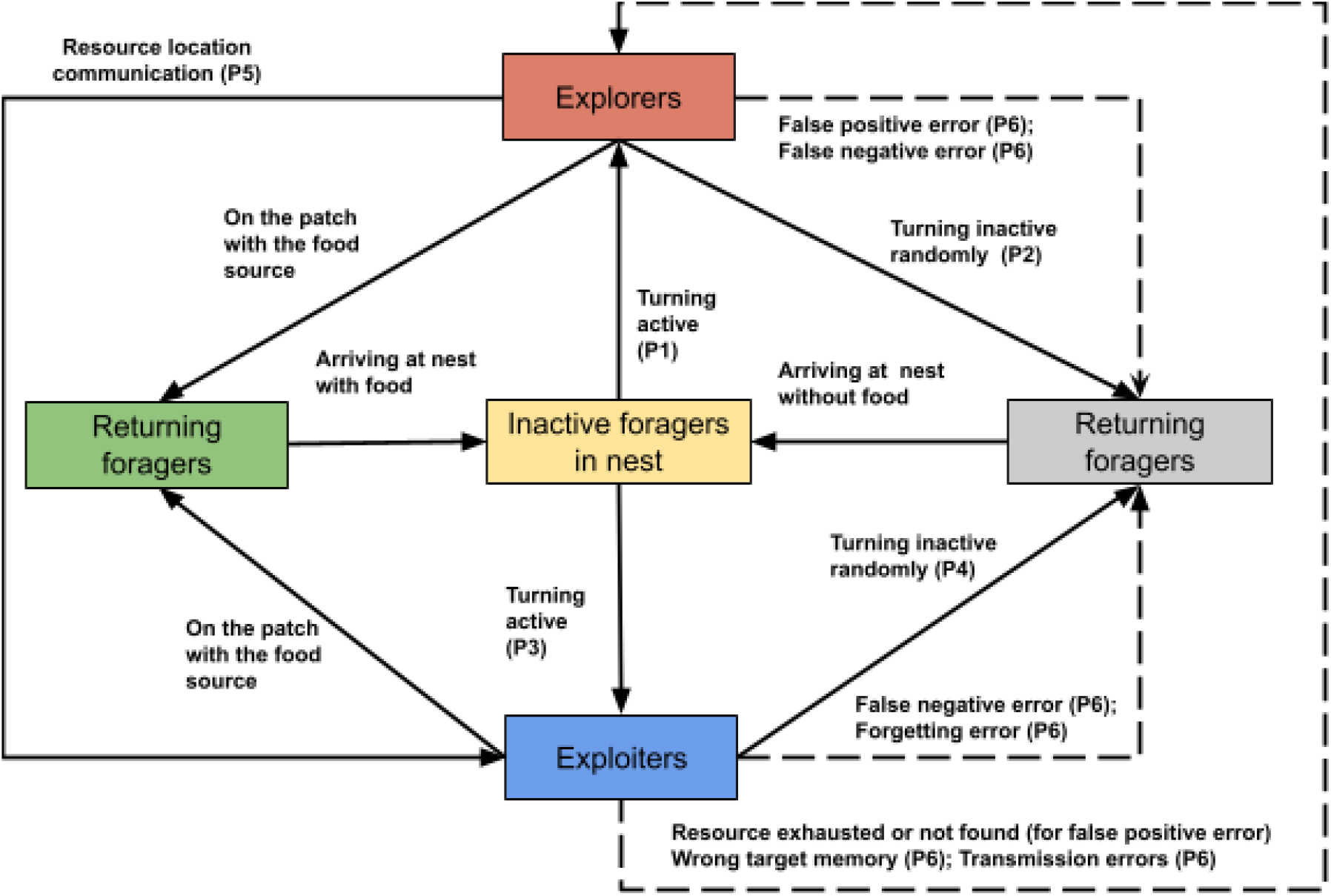
Behavior and states of agents (‘foragers’) in the simulation model. Possible state transitions are shown using solid lines. Foragers in the nest are inactive (yellow). If they have a resource location in their memory, they may leave and travel directly to this location (blue ‘exploiters’). If not, they may leave and search for a resource (red ‘explorers’). Once a resource is detected within their sensing radius, foragers remember this location, collect food from this resource, and return to the nest (green); upon arrival, they turn inactive (yellow) again. Traveling foragers (explorers or exploiters) may also probabilistically give up foraging and return to the nest (gray). Various error types may affect state transitions or memory content (shown using dotted lines). For more information on the transition probabilities, please see Table S1 in Supplementary file S1.

In summary, we found, as expected, that different error types acted in different parts of the foraging process. As a consequence however, we show that errors cannot be interpreted as favoring exploration over exploitation in general. Instead, when errors lead foragers to return to the nest earlier, they can severely curtail both exploration and foraging success, in part because in collective central-place foragers, a key aspect of exploration is to move outside of the area of high density of searchers around the nest itself. Those particular errors that lead foragers to attempt foraging in areas away from the nest thus allow the foragers and the colony to expand exploration and collect new information, potentially benefiting foraging success in some environments.

### Exploration vs. exploitation under errors and communication

Both the main benefit and the main cost of communication in collective foraging is that communication leads to targeted resource exploitation [99,103,106,113,114]. Since exploitation typically trades off with exploration (e.g., search can either be depth-first or breadth-first [9]), communication is therefore thought to imply decreased resource exploration [105,115–118]. It is thus surprising that we find increased exploration (as measured by resources discovered or area explored) with communication (Fig. 3), even without errors, likely primarily because communication drew foragers further away from the nest (panel i in Fig. S3 compared to Fig. 6 and Figs. S4-7 - note axis units). After communicating resources that were far from the nest became empty, this led foragers to start exploring at those distances, increasing overall exploration (both in terms of area and resources discovered). Without communication, searchers start from the nest itself, and thus are less likely to move into areas that have not yet been explored by other foragers from the same colony.

In our model, resource patches did not differ in their inherent quality, merely in their distance to the nest. Communication is thought to improve the ability of a colony to select the best resources [103], and see above), an aspect of performance we did not explicitly measure here. Colonies are still foraging selectively in the sense that they forage on resources closer to the nest than the average distance of resources to the nest in the environment (compare panels j and d in Fig. 6 and S3-7). However, we found that foraging close to the nest in general is associated with errors and generally poor overall foraging performance, because it is linked to less exploration and fewer resources discovered overall. Other than the fact that in our simulations resource quality was homogeneous, travel costs for food collection seemed to be minimal (compare time spent on collecting food, Fig. 4, to time spent searching for it, Fig. 3). While we sought to use empirically-validated parameter values (e.g., for the range over which colonies foraged and the time costs of travel), it would be interesting to quantify, for specific social insect species and their expected resource distributions [100,103], and how the time and energy costs of exploration compare to those of exploitation. This is likely to inform the tradeoff between exploration and exploitation specifically with regard to whether colonies benefit from exploring further afield even at the cost of relatively long travel times. If long travel is not particularly costly [106,119], then the evolution of error management and communication may be affected primarily by the benefit of leading foragers to make long trips to reduce overlap among searchers [120,121].

Overall, our model thus demonstrates that the amount of area explored is strongly dependent on processes that draw foragers away from the nest first, since areas close to the nest are likely to be explored by many individuals in any case. Errors may decrease movement ranges of foragers, reducing the area explored. This reframes the role of both errors and communication in the foraging process, as being strongly dependent on whether these factors act to promote effective exploration (e.g., far from the nest) in addition to whether they promote selection and exploitation of the highest quality resources.

### Pheromone-based communication did not perform well in our simulations

We implemented two pheromone-based communication types, namely ‘beacon’ and ‘trail’. Neither performed as well, with or without errors, as the direct, central-place, and blackboard communication types. Trail communication, on the other hand, generally did not produce outcomes that differed from ‘no communication’ very successful foraging outcomes (Fig. 3-4 and Fig. S3-S4). It is possible that this was a direct consequence of the small colony size modeled here (n=50), which can limit the successful build-up of a pheromone trail [76,122,123], Pheromone trails typically require not only build-up but also continuous reinforcement: in addition to the number of foragers, the number of trips made to the food source is thus critical.

Since we modeled food sources that were emptied after 5 trips, we thus may not have enabled sufficient repeat trips for foragers to generate and sustain successful trail recruitment (before the trail evaporates). The ‘beacon’ communication likely suffered from similar problems, but produced somewhat higher success, likely because it did draw foragers away from the nest (increasing area explored and search distance, comparing panels c and i in S11 with S8 and S12). In addition, the ‘beacon’ communication was less affected by the ‘forgetting’ error, likely because information was still contained in the environment even if foragers did not retain it in their memory.

### What do we learn from such a model

The purpose of models in biology, in general, is typically to demonstrate a general principle [93,94]. Individual-based simulations, like any numerical models, can only make statements about specific scenarios (that are defined by the parameter values and other assumptions used [66,87,124]. However, this can be particularly useful when demonstrating that particular processes or outcomes are *possible*, especially when the assumptions of the model match the system under study reasonably well. In addition, models often illustrate principles that are not difficult in hindsight but that were not apparent from verbal descriptions of a system [94]. Here, we modeled the behavioral rules and states of social insect foragers in a way that is identical or similar to other frequently used formal descriptions of them [66,125,126], and our specific parameter values are motivated by information from empirical studies (Table S1). What is new in our model is the inclusion of individual-level errors. We show that it is possible for such errors to disrupt the foraging process, and that such disruption, when the probability for error at the individual decision-point is equal, will depend on how frequently foragers find themselves at such decision points. This conclusion follows so directly that it should not require further justification; but it still was not obvious before the model demonstrated it (this is in fact often the benefit of models; e.g., see the famous case of Hardy- [94,127]).

Furthermore, our model demonstrates that in a complex system such as this one, non-intuitive and heterogeneous consequences of apparently similar processes are possible, in fact likely common. Specifically, we show that communication systems may interact with error types to produce results that differ both in magnitude and direction. Most importantly, we show that the common assumption that higher error levels generally equal more exploration and less exploitation is incorrect, and equally that the common assumption that communication always results in lower exploration is also incorrect; both of these principles are relevant broadly [9].

## Conclusion

Overall, using an agent-based modeling approach, we have shown that errors are detrimental to collective foraging and that communication can play an important role in mitigating the effects of errors. Importantly, our model illustrates that different error types may dramatically differ in their impact. A major driver of these differences seems to be that some errors simply have many more chances of occurring. We expect that sensory and cognitive systems evolve to prioritize minimizing such errors over others. Our model also demonstrates that not all errors are likely to increase exploration, and that spatial exploration in central place foragers depends on individuals having the chance and motivation to move far from the nest. Perhaps surprisingly, communication may increase rather than decrease spatial exploration for these reasons, because it draws foragers longer distances away from the nest, where they may search if initially exploited resources are emptied. These results suggest that the evolution of different communication systems in social insects may have been shaped not just by selection on improved exploitation of resources, but also by selection on exploration and error mitigation. Our model thus motivates further work, both empirical and theoretical, on the evolution of errors and error mitigation in individual and collective foraging.

## Materials and Methods

### Model overview

We developed an agent-based, spatially-explicit, continuous-space, discrete-time, stochastic model in NetLogo to simulate a social insect colony consisting of cooperating individuals (hereafter referred to as ‘foragers’). In the model, foragers collect resources from their environment using different communication types and are subject to different kinds of errors. Our model was implemented and run in NetLogo 6.3.0 [75] on multiple machines, including a high-performance computing cluster at the University of Arizona that uses a Linux-based command line interface. We analyzed more than 250,000 simulation runs in total. For addressing the first and second objectives, we varied error type and communication type, and for addressing third objective, along with these two parameters, we also varied the error probability per individual and timestep. All other parameters were held constant.

### Agents in the model

All simulations were run with 50 foragers and 100 resources (distributed in 10 clusters or 10 resources each, where each resource had a value of 5 trips worth of food); all parameters except for those of interest for the study were held constant at values judged to be broadly representative for social insect foraging (Table S1 in Supplementary file S1).

### The spatial and temporal scale of the model

We simulated a two-dimensional grid with a coordinate system consisting of 300 x 300 patches, where each patch was defined as being 10 forager body lengths wide (despite Netlogo’s framing of ‘patches’, the coordinate system is continuous). Therefore, the world allowed a foraging range of up to 1500 forager body lengths away from the central nest. Before each simulation run began, all the foragers were placed in the nest, and foragers returned regularly to the nest with or without food (Fig. 1). Each simulation was run for 2000 timesteps, with each step defined as equivalent to ten seconds (∼6 hrs of total simulated time per simulation).

### The foragers and their behavioral states, error types, and communication types

At any time, each forager was defined to be in one of five states as shown in Fig. 1. In addition to their behavioral state, foragers could also store one ‘target-patch’ in their memory - this is the location of the resource that they last visited. Foragers may update this ‘target-patch’ memory either by discovering or receiving information about a new resource location through communication. However, the memory is only updated if the new location is ‘better’, i.e., closer to the nest, than the location currently in memory (see Supplementary file S1 for more information).

Error types were modeled to affect how foragers detect, remember, and/or transmit the information on resource locations. Each simulation implemented one of five different individual-level error types (or no errors) - false positive errors (falsely sensing presence of a resource on an empty patch), false negative errors (failing to detect an active resource that could have been perceived), forgetting (forgetting the memory of a remembered resource location), wrong target memory (remembering a wrong resource location) and transmission errors (communicating a wrong resource location).

We use the term ‘communication type’ to refer to a type of information exchange characterized by which information is exchanged, and with what audience, irrespective of mechanistic details like signal modality. For example, some ants use pheromone trails to recruit nestmates to a food location; this type of communication essentially confers information about food location to a potentially unlimited number of receivers via a stigmergic signal, without requiring direct contact between individuals [76]. The honey bee waggle dance, on the other hand, also allows communication of food location, but only to a very limited number of receivers, who have to encounter the forager at a specific time and place [77]. We model communication like the waggle dance as ‘central place’ (exchange of information upon encounter of a forager and potential recruit at the nest), and pheromone trails as ‘trail’.

Communication in our model is implemented as receiver-initiated where the receiver extracts the ‘target-patch’ location from the memory of the sender. We implemented five different communication ‘types’, as well as ‘no communication’: direct communication (communicate anywhere in the world upon encounter), central-place communication (communicate at the nest upon encounter), blackboard communication (the nest stores information on the currently best resource, and workers extract information this information at the nest), beacon communication (workers deposit a signal at the food source that can be perceived from a distance), and trail communication (pheromone trail laid by workers who have discovered resources, of uniform strength in straight line to nest). In each simulation run, foragers used only one of the six communication types. Detailed information on the specifics of the error and communication types can be found in the Supplementary material file S1.

### Simulation process overview

Each simulation starts with a new randomly generated resource distribution and with all the foragers at the center of the map (the ‘nest’). Once the simulation starts, in each timestep, all foragers evaluate their behavior sequentially, potentially changing their state (Fig. 6), then communicate (if applicable, again independently and sequentially), then move (if applicable, again independently and sequentially). The patch variables and resource values are updated as and when the agents are on those patches or/and collect the resources. In the last timestep of each simulation run, a variety of output variables are calculated. We used NetLogo’s built-in BehaviorSpace function to run sets of simulations with varying inputs [75].

### Response variables quantified

We measured several outcomes of the collective foraging process: (1) overall foraging performance (total resource units collected), (2) exploration (resources discovered, resources discovered per unit area explored, area explored per unit time spent by foragers in searching, and Total forager-time spent searching), (3) exploitation (resource units collected per resource patch discovered, resources collected per unit time spent collecting resources, and time spent collecting resources per unit resource discovered), and (4) relative performances (resource units collected relative to the median resource units collected with no errors). See the ‘*Observation*’ section in Supplementary file S1 for more information.

### Statistical analyses

All statistical tests were performed in RStudio using R 4.5.2 [78]. To analyze the effect of different error and communication types on the foraging performance measures described above, we kept the *error-probability* value constant (N= 244,987, *error-probability* = 0.5), and used Kruskal-Wallis tests followed by post-hoc Dunn tests. The rest of the simulations were used for analyzing the effect of varying error probabilities on the number of resources collected. We fitted an inverted hill function (with parameters K and n) over the raw data over the whole range of error probability values. Other than base R, we used the following packages- *data.table* [79], *FSA* [80], *rcompanion* [81], *scales* [82]*, tidyverse* [83], and *viridis* [84].

## Supporting information

Supplementary materials (R code, model description and additional plots)

## Acknowledgements

We thank the University of Arizona HPC team, especially Sara Willis, for her help in streamlining the process of running jobs on the HPC clusters. We also thank Daniel Papaj, Judith Bronstein and Tanya Latty for their feedback on the manuscript.

