## Supplementary materials (R code, model description and additional plots) for "Frequent errors are the worst: robustness to individual failures in collective foraging": R code_2026.R

```
# Collective foraging & communication model in Netlogo -----------------
## (c) Anna Dornhaus 2020-2026
## Model by Anna Dornhaus, Kenny Chapin, Chinmay Joshi
## Github folder:
## https://github.com/dornhaus/CommunicationModel
# Here we analyze the impact of errors ------------
## This code is part B of a set of code files. 
### PART A: Import & data organization 
### PART B: Analysis and figures for paper on robustness (this file)
### PART C: Analysis and figures for paper on search
### PART D: Analysis and figures for paper on communication
## Outline of (this) code:
### I. Packages used
### II. Defining overarching parameters, e.g. graphics
### III. Data Import from local/github results folder 
### IV. Calculations
### V. Defining functions that generate figures
### VI. Figures in layout
############################################
# I. Packages used --------------------
library(rcompanion) # For compact letter labeling of Dunn's Test results
library(FSA) # For Dunn's test
library(tidyverse) # General data handling
library(data.table) # General data handling
library(scales) # for color scales
library(viridis) # for colors used
## Clean workspace ----------------------------
rm(list = ls(all.names = TRUE)) # Remove all objects in 'Environment' (variables, data, etc.)
gc() # Clear unused memory
# dev.off()
############################################
# II. Setting general options & paths [!!check this!!] ---------------
## Set the local directory for this repository
path_CommModel_repo <- "C:/Users/dornh/Dropbox/Github/CommunicationModel"
## Where to find the pre-sorted and formatted original data
sorted_results_path <- "C:/Users/dornh/Dropbox/RESEARCH/Netlogo Comm Model BKP & DATA/sorted_results 19_05_26"
figs_path <- "C:/Users/dornh/Dropbox/Github/CommunicationModel/Figures"

## Set this to FALSE if you want to keep all intermediate
## datasets and variables
variablecleanup <- TRUE  

## Save default graphics settings
opar <- par
############################################
# III. Import data from local results folder ----------------------
## Set working directory
setwd(paste(sorted_results_path, sep = ""))

## For this paper, we only use default parameters except we vary 
## communication mode and error type. For the last part of 'robustness shape'
## we also vary error probability. 
## We generated a table with just those data from all raw data in the import
## script.
complete_results <- as.data.frame(fread("err_x_comm_def.csv"))

############################################
# IV. Data formatting and calculations ------------------------------

## Ordering factor levels ---------------------------
### Fixed order
complete_results$MODULE_INDIV_ERRORS.incl <- factor(complete_results$MODULE_INDIV_ERRORS.incl, levels=c('none', 'false positive sensing', 'false negative sensing', 'wrong target memory', 'forgetting', 'transmission errors'))
complete_results$communication.mode.incl <- factor(complete_results$communication.mode.incl, levels=c('none', 'direct', 'central-place', 'blackboard', 'beacon', 'trail'))
levels(complete_results$MODULE_INDIV_ERRORS.incl) <- c('None', 'False positive sensing', 'False negative sensing', 'Wrong target memory', 'Forgetting', 'Transmission errors')
levels(complete_results$communication.mode.incl) <- c('None', 'Direct', 'Central-place', 'Blackboard', 'Beacon', 'Trail')
### Sort by performance (median resources collected)
complete_results$MODULE_INDIV_ERRORS.incl <- reorder(complete_results$MODULE_INDIV_ERRORS.incl, -complete_results$collected.resource.units, FUN = median)
complete_results$communication.mode.incl <- reorder(complete_results$communication.mode.incl, complete_results$collected.resource.units, FUN = median)
### But we want 'none' to be the first level no matter what
complete_results$MODULE_INDIV_ERRORS.incl <- factor(complete_results$MODULE_INDIV_ERRORS.incl, levels = c("None", setdiff(levels(complete_results$MODULE_INDIV_ERRORS.incl), "None")))
complete_results$communication.mode.incl <- factor(complete_results$communication.mode.incl, levels = c("None", setdiff(levels(complete_results$communication.mode.incl), "None")))
### Identify which level is now 'no errors' to be sure
pos_none <- which(levels(complete_results$MODULE_INDIV_ERRORS.incl)=="None")

## Defining colors, shapes, and transparency ---------------------------

#### Shapes: filled points for scatterplots except for 'none' as open circles
shape_list <- rep(19, nlevels(complete_results$MODULE_INDIV_ERRORS.incl))
shape_list[pos_none] <- 1

#### Standard colors for error levels
default_colorlist <- viridis(nlevels(complete_results$MODULE_INDIV_ERRORS.incl)-1, begin = 0.9, end = 0.1)
colorlist_errors_bl <- append("black", default_colorlist)
colorlist_errors_wh <- colorlist_errors_bl
colorlist_errors_wh[pos_none] <- "white" 

#### Standard colors for communication systems
colorlist_communication_bl <- magma(6, begin = 1, end = 0.3)
colorlist_communication_wh <- colorlist_communication_bl
colorlist_communication_wh[1] <- "white"

### Adjusting transparency for sample size to allow all error types to be equally visible
transparency_list_er <- 0.30/(0.30+0.001*as.vector(table(complete_results$MODULE_INDIV_ERRORS.incl)))

############################################
# V. Defining functions that generate figures ----------------------------------
## Function for better boxplot labeling - only needed here for the robustness ~ error prob plot
get_midpoint <- function(cut_label) {
  mean(as.numeric(unlist(strsplit(gsub("\\(|\\)|\\[|\\]", "", as.character(cut_label)), ","))))
}

## This is our standard plot showing the effect of different error types
error_fig <- function(outcome_numer, outcome_denomi=1, y_label="", y_max=-1, print_N=TRUE, print_effect=TRUE, format = "single_fig", results = complete_results) {
  if(format=="single_fig") {
    par(mfrow=c(2,3), oma = c(12,4,0,0))
    par(mgp=c(3, 1, 0), las=2, mar = c(1, 1, 2, 0.5))
    x_axis_on <- c(FALSE, FALSE, FALSE, TRUE, TRUE, TRUE)
    y_axis_on <- c(TRUE, FALSE, FALSE, TRUE, FALSE, FALSE)
  } else {
    if(format=="stacked") {
      # We're assuming this format: par(mfrow=c(6,1))
      par(mgp=c(3, 1, 0), las=2, mar = c(0.5, 1, 1, 0.5))
      x_axis_on <- c(FALSE, FALSE, FALSE, FALSE, FALSE, TRUE)
      y_axis_on <- c(TRUE, TRUE, TRUE, TRUE, TRUE, TRUE)
    } else {
      if(format=="double") {
        par(mgp=c(3, 1, 0), las=2, mar = c(1, 1, 3, 0.5))
        x_axis_on <- c(FALSE, FALSE, FALSE, TRUE, TRUE, TRUE)
        y_axis_on <- c(TRUE, FALSE, FALSE, TRUE, FALSE, FALSE)
      }
    }
  }
  kw_output <- data.frame(matrix(ncol = 5, nrow = 0))
  colnames(kw_output) <- c("Communication", "Response", "N", "Statistic", "Pvalue")
  for(comm_type_index in 1:6) {
    comm_type <- levels(results$communication.mode.incl)[comm_type_index]
    results_panel <- subset(results, communication.mode.incl == comm_type)
    if(outcome_denomi==1) {
      graph_data <- cbind(results_panel[outcome_numer], results_panel$MODULE_INDIV_ERRORS.incl)
    } else {
      graph_data <- cbind(results_panel[outcome_numer] / results_panel[outcome_denomi], results_panel$MODULE_INDIV_ERRORS.incl)
      graph_data <- na.omit(graph_data)
      graph_data <- graph_data[is.finite(graph_data[,1]), ]
    }
    if(y_max==-1) {y_max <- max(graph_data[,1])}
    Nice_Plot <- boxplot(graph_data[,1] ~ graph_data[,2]
                         , xlab = ""
                         , ylab = ""
                         , main = ifelse(format=="single_fig"||format=="double",comm_type,"")
                         , ylim=c(-(y_max*0.1), y_max)
                         , col = colorlist_errors_wh
                         , bty = "l"
                         , range = 0
                         , cex.axis = 1
                         , xaxt = ifelse(x_axis_on[comm_type_index], 's', 'n')
                         , yaxt = ifelse(y_axis_on[comm_type_index], 's', 'n')
                         , yaxp = c(0, y_max, 5)
    )
    if(print_N) {
      nbGroup <- nlevels(as.factor(Nice_Plot$names))
      text( 
        x=c(1:nbGroup), 
        y=y_max*ifelse(format=="single_fig", 1, 1.1),
        cex = 0.5,
        paste(Nice_Plot$n,sep=""),
        xpd = TRUE
      )
    }
    if(print_effect) {
      pos_none <- 1 #which(levels(results_panel$MODULE_INDIV_ERRORS.incl)=="None")
      relative_medians <- Nice_Plot$stats[3,]/Nice_Plot$stats[3,pos_none]
      relative_effect <- round((relative_medians-1) * 100, digits = 0)
      text( 
        x=c(1:nbGroup), 
        y=Nice_Plot$stats[5,]+y_max * 0.08,
        cex = 0.7,
        paste(relative_effect,"%",sep="")  
      )
    }
    # Stats
    # Kruskal Wallis test: results written to file
    kw <- kruskal.test(graph_data[,1] ~ graph_data[,2])
    kw_row <- list("Communication"=comm_type
                   , "Response"=paste(outcome_numer, "/", outcome_denomi)
                   , "N"=nrow(graph_data)
                   , "Statistic"=round(kw$statistic[[1]], 1)
                   , "Pvalue"=round(kw$p.value, 5))
    kw_output <- rbind(kw_output, kw_row)
    # Dunn's post hoc test for pairwise comparisons
    dunn_test_out <- dunnTest(graph_data[,1] ~ graph_data[,2])
    #Reordering the letters as per the order in the graph
    fac_pairs <- expand.grid(Group2 = levels(graph_data[,2]), Group1 = levels(graph_data[,2]))
    pairs_inorder <- paste0(fac_pairs$Group1, " - ", fac_pairs$Group2)
    dunn_test_out$res <- dunn_test_out$res[match(pairs_inorder, dunn_test_out$res$Comparison), ]
    dunn_test_out$res <- na.omit(dunn_test_out$res)
    #Extracting the significance letters for the Dunn test
    dunn_test_letters <- cldList(P.adj ~ Comparison
                                 , data = dunn_test_out$res
                                 , threshold = 0.05
                                 , remove.zero = FALSE 
                                 )
    text( 
      x=c(1:nbGroup), 
      y=y_max * -0.05,
      cex = 0.7,
      paste(dunn_test_letters$Letter,sep="")  
    )
  }
  if(format == "single_fig") {
    if(y_label=="") {
      mtext(text = paste(outcome_numer, "/", outcome_denomi, sep = ""), 
            side = 2, line = 2, outer = TRUE, las = 0)
    } else {
      mtext(text = y_label, 
            side = 2, line = 2, outer = TRUE, las = 0)
    }
    mtext(text = "Error type", 
          side = 1, line = 10, outer = TRUE, las = 0)
  }
  write.table(kw_output, file = paste("KW", outcome_numer, "_", outcome_denomi, ".txt", sep = ""))
}
## Standard plot showing different communication types
comm_fig <- function(outcome_numer, outcome_denomi=1, y_label="", y_max=1, print_N=TRUE, print_effect=TRUE, format = "single_fig", results = complete_results) {
  if(format=="single_fig") {
    par(mfrow=c(2,3), oma = c(12,4,0,0))
    par(mgp=c(3, 1, 0), las=2, mar = c(1, 1, 2, 0.5))
    x_axis_on <- c(FALSE, FALSE, FALSE, TRUE, TRUE, TRUE)
    y_axis_on <- c(TRUE, FALSE, FALSE, TRUE, FALSE, FALSE)
  } else {
    if(format=="stacked") {
      # We're assuming this format: par(mfrow=c(6,1))
      par(mgp=c(3, 1, 0), las=2, mar = c(0.5, 1, 1, 0.5))
      x_axis_on <- c(FALSE, FALSE, FALSE, FALSE, FALSE, TRUE)
      y_axis_on <- c(TRUE, TRUE, TRUE, TRUE, TRUE, TRUE)
    } else {
      if(format=="double") {
        par(mgp=c(3, 1, 0), las=2, mar = c(1, 1, 3, 0.5))
        x_axis_on <- c(FALSE, FALSE, FALSE, TRUE, TRUE, TRUE)
        y_axis_on <- c(TRUE, FALSE, FALSE, TRUE, FALSE, FALSE)
      }
    }
  }
  kw_output <- data.frame(matrix(ncol = 5, nrow = 0))
  colnames(kw_output) <- c("Error", "Response", "N", "Statistic", "Pvalue")
  for(err_type_index in 1:6) {
    error_type <- levels(results$MODULE_INDIV_ERRORS.incl)[err_type_index]
    results_panel <- subset(results, MODULE_INDIV_ERRORS.incl == error_type)
    if(outcome_denomi==1) {
      graph_data <- cbind(results_panel[outcome_numer], results_panel$communication.mode.incl)
    } else {
      graph_data <- cbind(results_panel[outcome_numer] / results_panel[outcome_denomi], results_panel$communication.mode.incl)
      graph_data <- na.omit(graph_data)
      graph_data <- graph_data[is.finite(graph_data[,1]), ]
    }
    Nice_Plot <- boxplot(graph_data[,1] ~ graph_data[,2]
                         , xlab = ""
                         , ylab = ""
                         , main = ifelse(format=="single_fig"||format=="double",error_type,"")
                         , ylim=c(-(y_max*0.1), y_max)
                         , col = colorlist_communication_wh
                         , bty = "l"
                         , range = 0
                         , cex.axis = 1
                         , xaxt = ifelse(x_axis_on[err_type_index], 's', 'n')
                         , yaxt = ifelse(y_axis_on[err_type_index], 's', 'n')
                         , yaxp = c(0, y_max, 5)

    )
    if(print_N) {
      nbGroup <- nlevels(as.factor(Nice_Plot$names))
      text( 
        x=c(1:nbGroup), 
        y=y_max*ifelse(format=="single_fig", 1, 1.1),
        cex = 0.5,
        paste(Nice_Plot$n,sep=""),
        xpd = TRUE
      )
    }
    if(print_effect) {
      pos_none <- 1 # which(levels(results_panel$communication.mode.incl)=="none")
      relative_medians <- Nice_Plot$stats[3,]/Nice_Plot$stats[3,pos_none]
      relative_effect <- round((relative_medians-1) * 100, digits = 0)
      text( 
        x=c(1:nbGroup), 
        y=Nice_Plot$stats[5,]+y_max * 0.08,
        cex = 0.7,
        paste(relative_effect,"%",sep="")  
      )
    }
    # Stats
    # Kruskal Wallis test: results written to file
    kw <- kruskal.test(graph_data[,1] ~ graph_data[,2])
    kw_row <- list("Error"=error_type
                   , "Response"=paste(outcome_numer, "/", outcome_denomi)
                   , "N"=nrow(graph_data)
                   , "Statistic"=round(kw$statistic[[1]], 1)
                   , "Pvalue"=round(kw$p.value, 5))
    kw_output <- rbind(kw_output, kw_row)
    
    levels(graph_data[,2]) <- gsub("-", "", levels(graph_data[,2])) 
    dunn_test_out <- dunnTest(graph_data[,1] ~ graph_data[,2])
    #Reordering the letters as per the order in the graph
    fac_pairs <- expand.grid(Group2 = levels(graph_data[,2]), Group1 = levels(graph_data[,2]))
    pairs_inorder <- paste0(fac_pairs$Group1, " - ", fac_pairs$Group2)
    dunn_test_out$res <- dunn_test_out$res[match(pairs_inorder, dunn_test_out$res$Comparison), ]
    dunn_test_out$res <- na.omit(dunn_test_out$res)
    #Extracting the significance letters for the Dunn test
    dunn_test_letters <- cldList(P.adj ~ Comparison
                                 , data = dunn_test_out$res
                                 , threshold = 0.05
                                 , remove.zero = FALSE 
    )
    text( 
      x=c(1:nbGroup), 
      y=y_max * -0.05,
      cex = 0.7,
      paste(dunn_test_letters$Letter,sep="")  
    )
  }
  if(format == "single_fig") {
    if(y_label=="") {
      mtext(text = paste(outcome_numer, "/", outcome_denomi, sep = ""), 
            side = 2, line = 2, outer = TRUE, las = 0)
    } else {
      mtext(text = y_label, 
            side = 2, line = 2, outer = TRUE, las = 0)
    }
    mtext(text = "Communication type", 
          side = 1, line = 10, outer = TRUE, las = 0)
  }
  write.table(kw_output, file = paste("KWbycomm", outcome_numer, "_", outcome_denomi, ".txt", sep = ""))
}

############################################
# VI. Figures in layout --------------------------------------
# Reminder, this is the format for the function that draws figures:
#error_fig(outcome_numer, outcome_denomi=1, y_label="", y_max=1, print_N=TRUE, print_effect=TRUE, format = "stacked") {
results <- complete_results
setwd(figs_path)
# Fig 2 Performance by error and communication -----------------
pdf(file = "Fig2 performance.pdf", width = 6, height = 10)
par(mfrow=c(5,3), oma=c(10,5,1,1))
error_fig("collected.resource.units", y_max = 350, format = "double")
plot.new()
plot.new()
plot.new()
comm_fig("collected.resource.units", y_max = 350, format = "double")
mtext(text = "Overall performance [resource units collected]", 
      side = 2, line = 3, outer = TRUE, las = 0)
mtext(text = "Error types", 
      side = 1, line = -29, outer = TRUE, las = 0)
mtext(text = "Communication types", 
      side = 1, line = 7, outer = TRUE, las = 0)
dev.off()

############################################
# Fig. 3 Robustness with increasing error ------------------------------
# Fitting an inverse Hill function to robustness ~ error prob
## New data import ----------------------
# This part of the script requires the bigger dataset including all error
# probabilities. 
setwd(paste(sorted_results_path, sep = ""))
## Importing the full data
results_main <- fread("err_x_comm_all_error_prob.csv")
setwd(figs_path)
# Need to make sure factors are appropriately ordered:
results_main$MODULE_INDIV_ERRORS.incl <- factor(results_main$MODULE_INDIV_ERRORS.incl, levels=c('none', 'false positive sensing', 'false negative sensing', 'wrong target memory', 'forgetting', 'transmission errors'))
results_main$communication.mode.incl <- factor(results_main$communication.mode.incl, levels=c('none', 'direct', 'central-place', 'blackboard', 'beacon', 'trail'))
levels(results_main$MODULE_INDIV_ERRORS.incl) <- c('None', 'False positive sensing', 'False negative sensing', 'Wrong target memory', 'Forgetting', 'Transmission errors')
levels(results_main$communication.mode.incl) <- c('None', 'Direct', 'Central-place', 'Blackboard', 'Beacon', 'Trail')
results_main$MODULE_INDIV_ERRORS.incl <- factor(results_main$MODULE_INDIV_ERRORS.incl, levels=levels(complete_results$MODULE_INDIV_ERRORS.incl))
results_main$communication.mode.incl <- factor(results_main$communication.mode.incl, levels=levels(complete_results$communication.mode.incl))

### We calculate the no error medians for each combination of communication and error type.
no_error <- subset(complete_results, complete_results$MODULE_INDIV_ERRORS.incl=="None")
median_table <- data.frame()
median_table <- no_error %>%
  group_by(communication.mode.incl) %>% 
  summarize(median.noerror.performance = median(collected.resource.units, na.rm = TRUE))

## Calculating impact of errors
# Now we look up & insert these no-error-medians for all performance measures that
# match the correct communication system (everything else should be the same).
results_main <- results_main %>%
  inner_join(median_table, by = c('communication.mode.incl' = 'communication.mode.incl'))
# So now we have the 'reference' of the no error median outcome for each individual
# simulation
results_main$robustness <- 1 - ((results_main$median.noerror.performance 
                                 - results_main$collected.resource.units) /
                                  results_main$median.noerror.performance)
## Setup -----------------------
pdf(file = "Fig3 robustness.pdf", width = 10, height = 10)
y_axis_top <- 1.5
y_axis_offset <- c(0, 0.05)
y_axis_offset_bymedians <- 0.0002
# Make a page with five graphs (one for each error type, not for no error)
par(mfrow=c(6,5))
# Make larger bottom margin to enable vertical tick labels
# Reminder: margins are defined in order bottom, left, top and right 
par(oma = c(4,4,3,0), mar = c(0.5, 0.5, 2, 0.5), mgp = c(3, 1, 0), las = 2)
no_of_bins <- 100
break_seq <- seq(from=0, to=1, length.out=no_of_bins+1)
results_main$new_bin <- cut(results_main$error.probability, breaks=break_seq)
# Make a new table with communication system, error type, and respective
# KA and n values (50% robustness and Hill coefficient~steepness)
robustness_shape <- data.frame(Comm = character(),
                               Error = character(),
                               KA = numeric(),
                               n = numeric(),
                               stringsAsFactors=FALSE)
starting_values_model_fitting <- data.frame(Comm = character(),
                                            Error = character(),
                                            medians_or_raw = character(),
                                            KA = numeric(),
                                            n = numeric(),
                                            stringsAsFactors=FALSE)
KAlist <- c(1, 0.1, 5, 0.01, 20, 15, 0.001, 100, 1000, 0.0001)
nlist <- c(1, 0.1, 5, 0.01, 20, 15, 0.001, 100, 1000, 0.0001)
maxfitattempts <- length(nlist) * length(KAlist)
fitwhat <- "raw"
## Figures & Fits, looping over communication & error types ------------------------
for (comm_index in 1:nlevels(results_main$communication.mode.incl)) {
  communication_type <- levels(results_main$communication.mode.incl)[comm_index]
  if(comm_index==nlevels(results_main$communication.mode.incl)) {
    last_row<-TRUE
  } else {last_row<-FALSE}
  print(communication_type)
  # Subsetting and binning results
  results <- subset(results_main, communication.mode.incl == communication_type)
  ## LOOPING OVER ERROR TYPES
  for (err_index in 2:nlevels(results_main$MODULE_INDIV_ERRORS.incl)) {
    # PLOT FOR EACH ERROR TYPE
    # Subsetting results
    tested_error_type <- levels(results_main$MODULE_INDIV_ERRORS.incl)[err_index]
    print(tested_error_type)
    subset_results <- subset(results, MODULE_INDIV_ERRORS.incl == tested_error_type)
    ### Plotting the boxplot of robustness
    Nice_Plot <- boxplot(robustness ~ new_bin
                         , data = subset_results
                         , xlab = ""
                         , ylab = ""
                         , main = ""
                         , ylim = c(-0.1, y_axis_top)
                         , xlim = c(1, no_of_bins)
                         , col = colorlist_errors_wh[[err_index]]
                         , bty = "l"
                         , range = 0
                         , lwd = 0.5 #line width
                         , yaxt = 'n'
                         , xaxt = 'n'
    )
    if(last_row)
      axis(1, at = c(1, (0.5*no_of_bins),no_of_bins), labels = c("0", "0.5", "1"))
    if(err_index==2) {
      axis(2, at = c(0,0.5,1), labels = c(0,0.5,1))
    }
    ## Lines for 1 = no impact of error
    ## 0.5 = cutoff for determining 50% effectiveness
    abline(h=1, lwd=1, col="blue", lty = "dashed")
    abline(h=0.5, lwd=1, col="grey",  lty = "dashed")
    
    ## NOW COMES THE MODEL FITTING!
    if(fitwhat=="medians") {
      ## Fitting a model just to the medians of the bins
      # Note that x is here just the index of the bin
      x <- c(1:no_of_bins)
      y <- Nice_Plot$stats[3,] # medians from boxplot directly
      # Uncomment to make sure these are the correct points:
      #points(y~x, col = "blue", cex = 3)
      # Fitting the function in a 'try' to catch errors (warnings really)
      # if the model doesn't converge 
      KAstart <- 0
      nstart <- 0
      fitattempt <- 0
      fittedInvHillmedians <- NULL
      while(is.null(fittedInvHillmedians) && fitattempt<maxfitattempts+1){
        fitattempt <- fitattempt + 1
        KAstart <- KAlist[(fitattempt %% length(nlist)) + 1]
        nstart <- nlist[(fitattempt %/% length(nlist)) + 1]
        print(fitattempt)
        print(paste("KA:", KAstart))
        print(paste("n:", nstart))
        try({fittedInvHillmedians<-
          nls(
            y ~ 1 - (x^n / (KA^n + x^n)),
            start = list(KA=KAstart, n=nstart),
            algorithm = "port",
            lower = list(KA=0, n=0)
          )}
          , silent = TRUE)
      }
      if (!is.null(fittedInvHillmedians)) {
        starting_values_model_fitting[nrow(starting_values_model_fitting)+1, ] <-
          c(communication_type, tested_error_type, "medians", KAstart, nstart)
        # Calculate x and y values to display on graph
        modelinput <- c(1:no_of_bins)
        predictedvalues <- 
          predict(fittedInvHillmedians,list(x=modelinput))
        #points(predictedvalues ~ modelinput, col = "purple")
        lines(predictedvalues ~ modelinput, lwd=2, col = "purple", lty = "solid")
      } else {
        starting_values_model_fitting[nrow(starting_values_model_fitting)+1, ] <-
          c(communication_type, tested_error_type, "medians", NA, NA)
      }
    }
    if(fitwhat=="raw") {
      ## Fitting an inverted Hill function to the raw data
      # Note that here x is on a different scale
      x <- subset_results$error.probability
      y <- subset_results$robustness
      # Fitting the function in a 'try' to catch errors (warnings really)
      # if the model doesn't converge 
      KAstart <- 0
      nstart <- 0
      fitattempt <- 0
      fittedInvHill <- NULL
      while(is.null(fittedInvHill) && fitattempt<maxfitattempts+1){
        fitattempt <- fitattempt + 1
        KAstart <- KAlist[(fitattempt %% length(nlist)) + 1]
        nstart <- nlist[(fitattempt %/% length(nlist)) + 1]
        print(fitattempt)
        print(paste("KA:", KAstart))
        print(paste("n:", nstart))
        try({fittedInvHill<-
          nls(
            y ~ 1 - (x^n / (KA^n + x^n)),
            start = list(KA=KAstart, n=nstart),
            algorithm = "port",
            lower = list(KA=0, n=0)
          )}
          , silent = TRUE)
      }
      if (!is.null(fittedInvHill)) {
        starting_values_model_fitting[nrow(starting_values_model_fitting)+1, ] <-
          c(communication_type, tested_error_type, "raw", KAstart, nstart)
        ## Calculate x and y values to display on graph
        # First, these are the x-values for the model (ie. error probabilities)
        modelinput <- unname(sapply(Nice_Plot$names[1:no_of_bins], get_midpoint))
        # Calculate matching y values
        predictedvalues <- 
          predict(fittedInvHill,list(x=modelinput))
        # Now we need those y values in the positions of the actual boxes
        # Hence the 'x values' are not in terms of error probability
        lines(predictedvalues ~ c(1:no_of_bins), lwd=2, col = "red", lty = "solid")
        ## Now we save the coefficients of the model on the raw data in a table
        robustness_shape[nrow(robustness_shape)+1, ] <- c(communication_type, tested_error_type,unname(coef(fittedInvHill))[1], unname(coef(fittedInvHill))[2])
      } else {
        starting_values_model_fitting[nrow(starting_values_model_fitting)+1, ] <-
          c(communication_type, tested_error_type, "raw", NA, NA)
      }
    }
    
    # Plot sample sizes below bars
    #text(
    #  x=c(1:no_of_bins), 
    #  y=-0.05 - y_axis_offset,
    #  cex = 0.6,
    #  paste(Nice_Plot$n)  
    #)
    if (err_index == 4) {
      mtext(paste("Communication type:", communication_type)
            , side = 3, las = 1, line = 0.5
            , outer = FALSE, cex = 0.8
      )
    }
    if (comm_index == 1) {
      mtext(paste(tested_error_type)
            , side = 3, las = 1, line = 3
            , outer = FALSE, cex = 0.7
            , font = 2
      )
    }
  } # End of the loop for plotting error types in one row
} # End of the loop for plotting each communication type as a row
# Axis labels ------------------
mtext("Error probability"
      , side = 1, las = 1, line = 2.5
      , outer = TRUE, cex = 1
)
print(paste("Error probability bins: ", Nice_Plot$names[1:no_of_bins], collapse = " "))
mtext(paste("Relative performance")
      , side = 2, las = 3, line = 2
      , outer = TRUE, cex = 0.8
)
dev.off()

#############################################
# Fig 4 Exploration ----------------------------------------------------------
# Exploration (res discovered) is efficient search(res/area) * avoidance(area/time) * investment in search(time)
pdf(file = "Fig4 exploration.pdf", width = 8, height = 10)
layout(matrix(c(1,2,3,4,5,6,7, # Comm. types
                8,8,8,8,8,8,8, # Y-Axis label 1
                9,10,11,12,13,14,15, # First set of 6 graphs
                16,16,16,16,16,16,16, # Y-Axis label 2
                17,18,19,20,21,22,23, # 2nd set of 6 graphs
                24,24,24,24,24,24,24, # Y-Axis label 3
                25,26,27,28,29,30,31, # 3rd set of 6 graphs
                32,32,32,32,32,32,32, # Y-Axis label 4
                33,34,35,36,37,38,39 # 4th set of 6 graphs
                )
              , 7, 9, byrow = F)
       , widths=c(0.5,0.5,1,0.5,1,0.5,1,0.5,1)
       , heights=c(0.5,1,1,1,1,1,1)) 
par(mgp=c(3, 1, 0), las=2, mar = c(0.5, 0.5, 0.5, 0.5), oma=c(12,0,0,0))
plot.new()
plot.new() #Vertical label 1
text(0.7,0.5, labels = levels(results$communication.mode.incl)[1]
     , cex = 1, xpd = TRUE, srt = 90)
plot.new() #Vertical label 2
text(0.7,0.5, labels = levels(results$communication.mode.incl)[2]
     , cex = 1, xpd = TRUE, srt = 90)
plot.new() #Vertical label 3
text(0.7,0.5, labels = levels(results$communication.mode.incl)[3]
     , cex = 1, xpd = TRUE, srt = 90)
plot.new() #Vertical label 4
text(0.7,0.5, labels = levels(results$communication.mode.incl)[4]
     , cex = 1, xpd = TRUE, srt = 90)
plot.new() #Vertical label 5
text(0.7,0.5, labels = levels(results$communication.mode.incl)[5]
     , cex = 1, xpd = TRUE, srt = 90)
plot.new() #Vertical label 6
text(0.7,0.5, labels = levels(results$communication.mode.incl)[6]
     , cex = 1, xpd = TRUE, srt = 90)
mtext("Communication types", side = 2, line = -1.5, outer=TRUE
      , cex = 1, xpd = TRUE, las=0)
plot.new() #Vertical axis label
text(0.2,0.5, labels = "Resources discovered"
     , cex = 1.5, xpd = TRUE, srt = 90)
plot.new()
text(0.5,0.5, labels = "Exploration =", cex = 1, xpd = TRUE)
error_fig("total.resources.discovered", y_max = 70, format = "stacked")
plot.new() #Vertical axis label
text(0.2,0.5, labels = "Resources discovered/Area explored"
     , cex = 1.5, xpd = TRUE, srt = 90)
plot.new()
text(0.5,0.5, labels = "Search  \nefficiency *", cex = 1, xpd = TRUE)
error_fig("total.resources.discovered", "proportion.patches.explored", y_max = 2000, format = "stacked")
plot.new() #Vertical label
text(0.2,0.5, labels = "Area explored/Forager-time searching"
     , cex = 1.5, xpd = TRUE, srt = 90)
plot.new()
text(0.5,0.5, labels = "Avoidance among  \nsearchers *", cex = 1, xpd = TRUE)
error_fig("proportion.patches.explored", "prop_time_searching", y_max = 0.8, format = "stacked")
plot.new() #Vertical axis label
text(0.2,0.5, labels = "Total forager-time spent searching"
     , cex = 1.5, xpd = TRUE, srt = 90)
plot.new()
text(0.5,0.5, labels = "Investment\nin search", cex = 1, xpd = TRUE)
error_fig("prop_time_searching", y_max = 0.7, format = "stacked")
mtext(text = "Error type", 
      side = 1, line = 10, outer = TRUE, las = 0)
dev.off()

# Fig 5 Exploitation --------------------------------------------------
pdf(file = "Fig5 exploitation.pdf", width = 8, height = 10)
# Exploitation(res collected/res disc) is efficiency exploiting(res coll/time coll) * investment in collection(time coll/res disc)
layout(matrix(c(1,2,3,4,5,6,7, # Comm. types
                8,8,8,8,8,8,8, # Y-Axis label 1
                9,10,11,12,13,14,15, # First set of 6 graphs
                16,16,16,16,16,16,16, # Y-Axis label 2
                17,18,19,20,21,22,23, # 2nd set of 6 graphs
                24,24,24,24,24,24,24, # Y-Axis label 3
                25,26,27,28,29,30,31 # 3rd set of 6 graphs
                )
              , 7, 7, byrow = F)
       , widths=c(0.5,0.5,1,0.5,1,0.5,1,0.5,1)
       , heights=c(0.5,1,1,1,1,1,1)) 
par(mgp=c(3, 1, 0), las=2, mar = c(0.5, 0.5, 0.5, 0.5), oma=c(12,0,0,0))
plot.new()
plot.new() #Vertical label 1
text(0.7,0.5, labels = levels(results$communication.mode.incl)[1], cex = 1, xpd = TRUE, srt = 90)
plot.new() #Vertical label 2
text(0.7,0.5, labels = levels(results$communication.mode.incl)[2], cex = 1, xpd = TRUE, srt = 90)
plot.new() #Vertical label 3
text(0.7,0.5, labels = levels(results$communication.mode.incl)[3], cex = 1, xpd = TRUE, srt = 90)
plot.new() #Vertical label 4
text(0.7,0.5, labels = levels(results$communication.mode.incl)[4], cex = 1, xpd = TRUE, srt = 90)
plot.new() #Vertical label 5
text(0.7,0.5, labels = levels(results$communication.mode.incl)[5], cex = 1, xpd = TRUE, srt = 90)
plot.new() #Vertical label 6
text(0.7,0.5, labels = levels(results$communication.mode.incl)[6], cex = 1, xpd = TRUE, srt = 90)
mtext("Communication types", side = 2, line = -1.5, outer=TRUE, cex = 1, xpd = TRUE, las=0)
plot.new() #Vertical axis label
text(0.2,0.5, labels = "Resource units collected/Resource patches discovered", cex = 1.5, xpd = TRUE, srt = 90)
plot.new()
text(0.5,0.5, labels = "Exploitation =", cex = 1, xpd = TRUE)
error_fig("collected.resource.units", "total.resources.discovered", y_max = 6, format = "stacked")
plot.new() #Vertical axis label
text(0.2,0.5, labels = "Resources collected/Time spent collecting", cex = 1.5, xpd = TRUE, srt = 90)
plot.new()
text(0.5,0.5, labels = "Collection  \nefficiency *", cex = 1, xpd = TRUE)
error_fig("collected.resource.units", "prop_time_collecting", y_max = 3000, format = "stacked")
plot.new() #Vertical axis label
text(0.2,0.5, labels = "Time spent collecting/Resources discovered", cex = 1.5, xpd = TRUE, srt = 90)
plot.new()
text(0.5,0.5, labels = "Investment\nin exploitation", cex = 1, xpd = TRUE)
error_fig("prop_time_collecting", "total.resources.discovered", y_max = 0.04, format = "stacked")
mtext(text = "Error type", 
      side = 1, line = 10, outer = TRUE, las = 0)
dev.off()

###################################
# Fig. 6 What causes performance? ----------------------------
## Setup
results <- complete_results
setwd(figs_path)
y_max <- 200
reg_lines <- FALSE
## Define basic scatterplot
xy_plot <- function(results_fig, xfactor, xlabel = "", yax = TRUE, panel_letter = "a") {
  outcome <- results_fig[xfactor]
  plot(collected.resource.units ~ outcome[,1]
       , data = results_fig
       , col = alpha(colorlist_errors_bl[results_fig$MODULE_INDIV_ERRORS.incl], transparency_list_er[results_fig$MODULE_INDIV_ERRORS.incl])
       , pch = shape_list[results_fig$MODULE_INDIV_ERRORS.incl]
       , ylim = c(0, y_max)
       , ylab = ""
       , xlab = ifelse(xlabel=="", xfactor, xlabel)
       , yaxt = ifelse(yax, 's', 'n')
  )
  text(min(outcome[,1], na.rm = TRUE),y_max*0.95, pos = 4, labels = panel_letter, col = "black", font = 2, cex = 2)
  if(reg_lines)
  for(err_index in 1:6) {
    err_type <- levels(results_fig$MODULE_INDIV_ERRORS.incl)[err_index]
    reg_line_dat <- subset(results_fig, results_fig$MODULE_INDIV_ERRORS.incl==err_type)
    x <- reg_line_dat[xfactor][,1]
    y <- reg_line_dat$collected.resource.units
    abline(lm(y ~ x), col = colorlist_errors_bl[err_index], lwd = 2)
  }
}
## Define basic horizontal boxplot
margin_boxpl <- function(results_fig, xfactor) {
  response <- results_fig[xfactor]
  boxplot(response[,1] ~ MODULE_INDIV_ERRORS.incl
          , data = results_fig
          , xaxt = 'n'
          , yaxt = 'n'
          , ylab = ""
          , xlab = ""
          , frame = FALSE
          , range = 0
          , col = colorlist_errors_wh
          , horizontal=TRUE
  )
}
## Loop over communication systems to make one figure each
for(comm_type_index in 1:6) {
  comm_type <- levels(results$communication.mode.incl)[comm_type_index]
  results_fig <- subset(results, communication.mode.incl == comm_type)
  pdf(file = paste("Fig6_", comm_type, "_scatterplots.pdf"), width = 10, height = 6)
### Layout: 4 rows, 6 columns, but the first and third row are small, for boxplots
  layout(matrix(c(1,2,3,4,5,6,7,8,9,10,11,12,13,14,15,16,17,18,19,20,21,22,23,0), 4, 6, byrow = T)
         , widths=c(1.2,1,1,1,1,1), heights=c(5,1,5,1)) 
  par(oma=c(0,2,0,0), mgp=c(2,1,0))
### Row with scatterplots
  par(mar=c(3,3,0.5,0.5))
  xy_plot(results_fig, "prop_time_searching", xlabel = "Prop. time searching", panel_letter = "a")
  par(mar=c(3,0.5,0.5,0.5))
  xy_plot(results_fig, "timesteps.firstcollected", xlabel = "Time to first discovery", yax = FALSE, panel_letter = "b") 
  xy_plot(results_fig, "proportion.patches.explored", xlabel = "Area explored", yax = FALSE, panel_letter = "c") 
  xy_plot(results_fig, "average.resource.distance", xlabel = "Avg. resource distance", yax = FALSE, panel_letter = "d") 
  xy_plot(results_fig, "total.trips", xlabel = "Total trips", yax = FALSE, panel_letter = "e") 
  xy_plot(results_fig, "efficiency", xlabel = "Efficiency", yax = FALSE, panel_letter = "f") 
  legend("topright", legend = levels(results_fig$MODULE_INDIV_ERRORS.incl), col = colorlist_errors_wh, pch = shape_list, cex = 0.8)
### Row with boxplots
  par(mar = c(1,3,0,0.5)) # bottom, left, top, right
  margin_boxpl(results_fig, "prop_time_searching")
  par(mar = c(1,0.5,0,0.5)) # bottom, left, top, right
  margin_boxpl(results_fig, "timesteps.firstcollected")
  margin_boxpl(results_fig, "proportion.patches.explored")
  margin_boxpl(results_fig, "average.resource.distance")
  margin_boxpl(results_fig, "total.trips")
  margin_boxpl(results_fig, "efficiency")
### Second row with scatterplots
  par(mar=c(3,3,0.5,0.5))
  xy_plot(results_fig, "prop_time_collecting", xlabel = "Prop. time collecting", yax = FALSE, panel_letter = "g") 
  par(mar=c(3,0.5,0.5,0.5))
  xy_plot(results_fig, "total.resources.discovered", xlabel = "# res. discovered", yax = FALSE, panel_letter = "h") 
  xy_plot(results_fig, "average.search.distance", xlabel = "Avg. search dist.", yax = FALSE, panel_letter = "i") 
  xy_plot(results_fig, "average.exploited.resource.distance", xlabel = "Avg. exploited res. dist.", yax = FALSE, panel_letter = "j") 
  xy_plot(results_fig, "unsuccessful.search.trips", xlabel = "Unsuccessful search trips", yax = FALSE, panel_letter = "k") 
### Final panel is a boxplot
  boxplot(collected.resource.units ~ MODULE_INDIV_ERRORS.incl
          , data = results
          , col = colorlist_errors_wh
          , ylim = c(0, y_max)
          , ylab = ""
          , yaxt = 'n'
          , xaxt = 'n'
          , xlab = "Error types"
          , frame = FALSE
          , range = 0
          , horizontal=FALSE
  )
  text(0,y_max*0.95, pos = 4, labels = "l", col = "black", font = 2, cex = 2)
### Second row with boxplots
  par(mar = c(1,3,0,0.5)) # bottom, left, top, right
  margin_boxpl(results_fig, "prop_time_collecting")
  par(mar = c(1,0.5,0,0.5)) # bottom, left, top, right
  margin_boxpl(results_fig, "total.resources.discovered")
  margin_boxpl(results_fig, "average.search.distance")
  margin_boxpl(results_fig, "average.exploited.resource.distance")
  margin_boxpl(results_fig, "unsuccessful.search.trips")
### Outer margin label
  mtext(paste("Resources collected with communication type: ", comm_type), side = 2, line = 0, outer = TRUE)
  dev.off()
}

############################
# Fig. S1 Relationship in time investment -----------------------
xy_plot_flexy <- function(results_fig, xfactor, yfactor, xlabel = "", ylabel = "", yax = TRUE, panel_letter = "a", reg_lines = FALSE, y_max=-1) {
  predictor <- results_fig[xfactor]
  response <- results_fig[yfactor]
  if(y_max==-1) {y_max <- max(response[,1])}
  plot(response[,1] ~ predictor[,1]
       , col = alpha(colorlist_errors_bl[results_fig$MODULE_INDIV_ERRORS.incl], transparency_list_er[results_fig$MODULE_INDIV_ERRORS.incl])
       , pch = shape_list[results_fig$MODULE_INDIV_ERRORS.incl]
       , ylim = c(0, y_max)
       , xlim = c(0, max(predictor[,1]))
       , ylab = ifelse(ylabel=="", yfactor, ylabel)
       , xlab = ifelse(xlabel=="", xfactor, xlabel)
       , yaxt = ifelse(yax, 's', 'n')
  )
  text(min(predictor[,1], na.rm = TRUE),y_max*0.95, pos = 4, labels = panel_letter, col = "black", font = 2, cex = 2)
  if(reg_lines)
    for(err_index in 1:6) {
      err_type <- levels(results_fig$MODULE_INDIV_ERRORS.incl)[err_index]
      reg_line_dat <- subset(results_fig, results_fig$MODULE_INDIV_ERRORS.incl==err_type)
      x <- reg_line_dat[xfactor][,1]
      y <- reg_line_dat[yfactor][,1]
      if(length(x)>0) {
        abline(lm(y ~ x), col = colorlist_errors_bl[err_index], lwd = 3, lty = 2)
      }
    }
}
results <- subset(complete_results, complete_results$communication.mode.incl=="Central-place")
par(mfrow=c(1,1), oma=c(0,0,0,0))
par(mar=c(5,5,0.5,0.5))
xy_plot_flexy(results, "prop_time_searching", "prop_time_collecting", y_max = 0.2 , panel_letter = "", reg_lines = TRUE, xlabel = "Prop. time searching", ylabel = "Prop. time spent collecting")
legend("topright", legend = levels(results$MODULE_INDIV_ERRORS.incl), col = colorlist_errors_bl, pch = shape_list)
# Fig. S2 Prop time inactive ---------------------------
error_fig("prop_time_inactive", y_label = "Prop time inactive", y_max= 0.68)

#############################################################################
```
