## Supplementary materials (R code, model description and additional plots) for "Frequent errors are the worst: robustness to individual failures in collective foraging": Supplementary file S1_2026.pdf

Table S1: Parameter table. Detailed description of the numerical parameters that were included in the model.

| **Variables** | **Description** | **Default values** | **Source** |
|---|---|---|---|
| patch size | Basic space | 10 forager body length | Social insect forager body sizes are far smaller than the distances that they travel for foraging. So, to ensure that the patches are much larger than individual foragers, we choose this as the patch length (see under the '*world-size*' for more information) [1,2]. |
| timestep | Basic time unit | 1 tick = 10 seconds | This captures the minimum time scale we need to capture different steps during social insect foraging such as exploration, loading the food at the food source, unloading the food at the nest, and returning to the food source until it is exhausted (See below under *simulation-duration* for more information). For example, honey bee foragers may a few seconds to around 10 minutes |

| | | | |
|---|---|---|---|
| | | | to unload the nectar |
| resource unit | The amount of food that can be carried by a forager during one trip | 1 unit | One forager load is the minimum resource unit of interest in our model. This may be the equivalent of one seed or prey item, or several µl of nectar [3–5] . |
| simulation-duration | Duration of a simulation run | 2000 ticks | We aimed to capture the typical foraging duration in a day, based on the idea that new resources have to be discovered on the next day. Active foraging time per day ranges from 6 to 24 hours with considerable variation within and across social insect species. Here, 1 tick is 10 seconds, so 2000 ticks is 5.5 hours [6–8] . |
| number-of-ants | Colony size | 50 | Social insect colony sizes range from 1 individual to more than a million workers per colony. However, most species have mature colonies around 100, and only 5-30% of the workers typically |

| | | | |
|---|---|---|---|
| | | | participate in foraging [9]. |
| world-size | Total size of the world available to the foragers | 300 x 300 patches | Most honey bee foraging trips are within 1.6 km, and most ant trips are within 5-7 m. So to capture most of these trips, we chose the default size to be 300 x 300 (3000 x 3000 body lengths) [10–12]. |
| cluster-radius | Size of a resource cluster (with multiple resource units) | 5 | A single resource patch with multiple individual resource units (flower patch, for example) can range from 0.2 $m^2$ to about 445 $m^2$ (equivalent to ranging from 5 patches to 700 patches) [13]. Foragers collect seeds that are distributed over a large patch with 1 to 50 seeds per centimeter square (equivalent to 1 to 150 seeds per patch here) [14,15]. |
| number-of-clusters | Total number of resource clusters available in the world | 10 | There is a large variation in the number of resource clusters of a given resource. For example, a flower patch can have as low as 10 flowers |

| | | | |
|---|---|---|---|
| | | | to as high as 5000 flowers [16,17]; Seed densities can range from 1 seed to 50 seed per centimeter square (equivalent of 1 to 150 seeds per patch here) [14,15]. |
| resources-per-clusters | Total number of resource units in each cluster | 10 resource units | There is considerable variation in the number of resources per cluster. For example, Johnson and Hubell (1975) found that *Casia* plants have 20 to 100 flowers per cluster within the same area [15,16,18,19]. |
| value-per-resource | Value of each resource unit within a cluster | 5 | Social insects like ants often recruit multiple workers to carry a prey item back to the nest (e.g., 30-60 workers in [20]). Pollinators like honey bees or bumble bees may need to make multiple trips to empty flowers on a given plant (crop size- 50 microliters [21] and total nectar volume across all flowers on a plant can exceed the crop size of an individual bee [22,23]). So, to |

| | | | |
|---|---|---|---|
| | | | not have resources emptied in a single trip, we choose the default value of 5 (i.e., 5 trips needed to empty the resource). |
| probability-activate-exploiters | Probability of inactive forager turning into an exploiter (with memory) | 0.1 / time step | The default value of 0.1 means that inactive foragers with memory of resource location, on average, will become active after 10 time steps. This accounts for the time spent by foragers going inside the nest, unloading their crop contents to other foragers via trophallaxis, interacting with other workers in the nest, and then making their way out to the nest. For example, honey bees spend a few seconds to around 10 minutes within the hive transferring the nectar to other nestmates via trophallaxis [24–26]. |
| probability-activate-explorers | Probability of inactive forager turning into an explorer (no | 0.0037 / time step | A default value of 0.0037 implies that inactive workers will become active after an expected time of 300 |

| | | | |
|---|---|---|---|
| | memory) | | timesteps (i.e., 50 min). This default value is the same as the inactivation probability for explorers, which makes it equally likely to be either active or inactive. In addition to this, frequent inactivation may allow explorers in the nest to extract information from the inactive exploiters in the nest. For example, honey bee foragers may spend a long time inside the nest (~1 hour) waiting to get recruited by other foragers via waggle dance [24,27,28]. |
| probability-inactivate-exploiters | Probability of an exploiter turning inactive (returning to nest) | 0.001 / time step | A default value of 0.001 implies that a forager with memory of a resource, on average, will spend 1000 timesteps (~2.7 hours) before it turns inactive. This is based on the idea that even successful foragers will not continue foraging indefinitely. For example, foragers may give up foragers within a few |

| | | | |
|---|---|---|---|
| | | | minutes to an hour [7,29,30]. |
| probability-inactivate-explorers | Probability of an explorer turning inactive (returning to nest) | 0.0037 / time step | A default value of 0.0037 means that a forager searching for resources, if unsuccessful, will eventually become inactive (here after an average of 50 minutes). For example, honey bee foragers give up foraging after searching for a few minutes to an hour [7,29,30]. |
| prop-starting-yellow | Proportion of individuals that are inactive at the start of the simulation | 0.25 | To prevent artefacts from all foragers potentially leaving the colony simultaneously (and returning after a similar time as well), we initially set a proportion of foragers as ‘inactive’. These become active over time with the ‘probability-activate-explorers /exploiters’ probabilities per timestep. |

| | | | |
|---|---|---|---|
| tortuosity | Standard deviation of turning angle of explorers | 60 degrees | We model search as a simple, correlated random walk with normally distributed turning angles around a mean of 0. The actual 'tortuosity' of exploring social insect foragers may vary widely and adaptively. The level of tortuosity modeled here (defined by the turning angle distribution) was chosen to match the temporal and spatial scale of foraging modeled, such that workers were likely to explore a good portion of the map but not run into the edges of the map very often [31–33]. |
| step-length | Forager traveling speed | 1 patch/time step | Social insect foragers travel at different speeds by walking (ants: e.g. ~0.4 m/s -> ~140 body lengths/s, -> 1.4 patches/time step) or by flying (bees, wasps: e.g. ~20 m/s -> ~2000 body lengths/s -> ~20 patches/time step) [32,34–36]. |

| | | | |
|---|---|---|---|
| sense-radius | How far the forager can detect the presence of a resource, another forager, a pheromone, or a patch. | 5 forager body lengths (0.5 patches) | Social insect foragers can detect food and the presence of other foragers using different modalities, and the distance of detection changes with the type of modality. For example, Carpenter ants [37] and Argentine ants can detect the presence of a nestmate and pheromone trail respectively within 1cm [38]. |
| error-level | How far the forager deviates from the original location (only applicable for transmission and wrong target memory errors) | 20 patches | For transmission and wrong memory errors, a novel 'faulty' location had to be selected that would either be transmitted or remembered respectively. The coordinates of this faulty location come from a normal distribution with the mean value equal to the coordinates of the original location and standard deviation equal to the 'error-level.' This was to ensure that the faulty target would not be arbitrarily far away from the nest, but far |

| | | | |
|---|---|---|---|
| | | | enough from the actual target so that the forager would not immediately encounter the same resource again. |
| error-probability | The probability that an individual forager makes an error | 0.5 / applicable time step | We chose a probably unrealistically high error probability to be able to clearly illustrate the effects of errors. |
| signal-strength | Probability that a forager will extract the signal from another forager when both are in each other's sense radius | 0.1 / applicable time step | The probability of foragers communicating presence of food or reinforcing the existing signal leading towards the food can vary. For example, 39-44% *Lasius niger* foragers reinforce the pheromone trail [39]. Probability of a honey bee forager following another forager's waggle dance varies between 0.2-0.9 [40]. |
| pheromone-deposited | Amount of pheromone deposited in total | 1000 | We adjusted this value in combination with the pheromone-fade-rate, see below. |
| pheromone-fade-rate | The rate at which the | 0.01 | Here, we considered the |

| | | | |
|---|---|---|---|
| | pheromone disappears | | short-term pheromone trail used for recruiting workers to retrieve an ephemeral food source. Such trails need continuous reinforcement from the workers, and are typically very short-lasting compared to the average foraging duration of the whole colony (10-45 minutes, which is equivalent to 60 to 270 time steps here) [41–44]. |

### Detailed model description

The model description is formatted as per the ODD (Overview, Design concepts, Details) format for individual- and agent-based models [45–47].

#### Model purpose

The purpose of our model was to investigate the effect of different individual-level errors on collective foraging in social insects, and whether and how different communication types affect performance robustness against these errors. We aimed to contribute to a broad, qualitative understanding of the effect of individual errors on collective performance. However, we also aimed to situate our model in the region of applicable parameter space that generally reflects the constraints and conditions under which social insects search for and exploit food resources. We therefore chose parameter values that represent foraging in a general social insect colony (see Table S1 for empirical references). Nonetheless, our model must be considered an abstracted, simplified model of social insect foraging. In addition, we identified different types of performance measures which may be relevant in different social insects, and which are not all equally susceptible to the effect of different error types (See 'response variables quantified' section in the main manuscript).

#### Entities, state variables, and scales

Formally, there are three types of agents in this model - the 'patches', i.e., the grid cells in the world map, the resources exploited by the foragers (i.e., food sources), and the individual foragers.

*The 'world', 'patches', and spatial scale*

The NetLogo world consists of a two-dimensional grid of discrete patches or coordinates. We used a coordinate system of 300x300 of these discrete patches. We chose the parameter values based on an interpretation that a patch has a side length of 10 forager body lengths. Thus, the simulated world allows foragers to explore at distances of up to 1500 times their body length from the nest (Table 1). The patches themselves are treated as agents that store information such as whether they have been explored (i.e., visited by any foragers), and how much pheromone has been deposited on them (if foragers are using beacon or trail communication). The patch at the origin (center) of the coordinate system is named 'the nest'. All foragers are at the nest before a simulation run begins (see under 'Simulation process overview' for more details). We interpret one time step to be equivalent to ten seconds. Since we ran simulations for 2000 timesteps, this can be seen as equalling ~ 6 hours. This total simulation duration is based on the assumption that typical resource location memory, communicated information, and foraging trips build up and are relevant during a single day, but essentially reset on the following day. In fact, foragers of ant species in *Solenopsis, Pheidole, Temnothorax, Lasius,* and other genera forage on resources that are ephemeral (insect carrion or fallen fruits) [48]. Similarly, social insects like honeybees or harvester ants tend to discover and forage on new patches or consistently visit the same patch of flowers [49,50] or seeds [5,6] each day, although they can remember patch locations for longer if patches persist [51,52]. Thus, our conclusions may be particularly relevant for species foraging on fast-changing (on the scale of a day) resource environments. However, we argue that many conclusions are likely to be general and apply to other timescales (see 'Discussion' section).

*The resources*

The resources are generated once at the beginning of the simulation and do not refill over time (i.e., they are assigned a 'value' and position once during each simulation run). Each visit by a forager removes one unit of 'value', until the 'value' reaches zero. The resource is then considered empty, and foragers no longer react to it. Our model thus emulates the challenge of initial discovery and quick exploitation of a single set of resources, such as nectar-filled flowers early in the morning when bees start to forage [7], or seed-rich patches sought by harvester ants [6], before a fixed deadline (e.g., sunset in bees when flight is no longer possible, or when daytime temperatures get too hot for foraging in harvester ants). Social insects face competition from other colonies nearby (both inter- and intraspecific), which can further reduce the time they can invest in foraging [53–56]. In this study, we do not examine all possible resource environments; instead, we focus on the common situation of somewhat patchily distributed resources (as would be the case for the mentioned honey bees or harvester ants, see Table 1). This patchy resource distribution was achieved by placing ten resource clusters (with ten identical resource patches per cluster (resource value of five per unit)) in a uniform random distribution. Each cluster had a fixed radius of 5 patches with resources randomly distributed within each cluster for each simulation run.

*The foragers, error types, and behavioral states*

*Foragers and their behavioral states*

The foragers are the agents that move around the NetLogo world actively throughout the simulation duration to explore and collect resources. At any given timestep in a simulation run, each forager is in one of the five states below which are denoted in our model as 'colors'; (See

Fig. 1). In addition to their state ('color'), foragers also store a '*target-patch*' in their memory. This is the position of the last exploited resource, or alternatively, is information about a position that was communicated by another forager. Foragers (and the nest) can remember and communicate *target-patch* information to other foragers in their *sense-radius* (See the 'Interaction' section below for details on communication types in our model).

- Inactive (yellow) - All foragers that return to the nest (with or without a resource) turn inactive. These foragers may then turn to being an explorer or an exploiter, depending on whether they have a memory of a resource location with an activation probability.
- Explorers (red) - Explorers are the ones that are exploring their environment, i.e. searching for resources. These foragers are leaving or are outside the nest, and therefore do not have a resource location in their memory. Exploiters may become explorers if they forget the resource location, or arrive at an empty or nonexistent resource location due to errors.
- Exploiters (blue) - Exploiters are the ones that are traveling to a known resource location (*target-patch*). Inactive foragers with a resource location in memory become exploiters with an activation probability, but also if they receive location information from another forager. Any explorers can directly turn into exploiters if they receive such information.
- Returning with resource (green) -Any forager that finds a resource travels back to the nest in a straight line; once they arrive at the nest, they turn inactive.
- Returning without resource (gray) - Explorer and exploiters turn inactive probabilistically with an inactivation probability (or when errors occur, see below) and return back to the nest without any resource (in a straight line). Once at the nest, they turn inactive.

*Error types and how and when they come into play*

Different error types can affect how foragers detect, remember, and/or transmit the target-patch information. Each simulation modeled one of five different individual-level errors (or no errors).

a. No error: Foragers remember and communicate exact resource locations.
b. False positive errors: Foragers (only explorers) may falsely sense the presence of a resource on an empty patch; they immediately return to the nest (as if having just collected food). As a consequence, they commit to memory and may communicate a location that lacks a resource. For example, bumble bees can mistakenly visit unrewarding flowers that mimic rewarding flower colors [57].
c. False negative errors: Foragers (both explorers and exploiters with memory) may fail to detect an active resource (i.e., either when searching for a new resource or attempting to return to a known resource) in their *sense-radius*, and immediately return to the nest. Any memory of a resource location remains, and therefore, exploiters may make further attempts to find the known resource on their next trip. For explorers, this will typically lead to a continued search for (other) resources after they turn active. For example, smaller bumble bees have much lower optical resolution in their visual system, and may miss small or far-away flowers [58].
d. Forgetting errors: Exploiters with memory of a known food location may forget their remembered resource location. They turn inactive and return to the nest, and turn into explorers (i.e., no memory) when they become active again. For example, *Cataglyphis niger* ants forgot the turns in a maze that lead to a food source over a two-week period [59].

e. Transmission errors: Explorers extract location information from an exploiter within its *sense-radius* that is in fact not the *target-patch* of the exploiter (i.e., a non-resource patch). The specific implementation of this error type somewhat varies by communication types. However, for all the communication types, the signal receiver ends up going to an empty location, while the signal sender continues going to the correct location (i.e., its *target-patch*). In communication types where a location memory is directly copied to a receiver ('direct', 'central-place', and 'blackboard', see below), the received coordinates (both x and y coordinates) are changed to random coordinates. These random coordinates come from a normal distribution with the mean value equal to the coordinates of the original location and standard deviation equal to the *'error-level'* (see Table 1). In pheromone communication types ('beacon' and 'trail'), the forager, rather than following the pheromone trail, goes in a random direction (and thus continues to explore for resources). This random direction comes from a normal distribution centered around zero with a standard deviation equal to the *'error-level'*. For example, the honey bee waggle dance has an inherent angular error that decreases with increasing distance of the food source (that is being communicated by the dance) from the nest [60].

f. Wrong target memory: Exploiters remember coordinates of a non-resource location instead of the actual resource coordinates. These changed coordinates are random numbers that are chosen from a normal distribution with a mean value equal to the coordinate of the original location (for both x and y coordinates), and the standard deviation equal to the 'error-level.' Their current food collection trip is not impacted, but this implies they may communicate and return on the next trip to an empty location. For example: Bumble bees exhibit preferences towards resources that have attributes that are a combination of two independently rewarding resources (merging of memories to create another memory that they have not experienced and therefore may not exist) [61].

### Process overview

Every new simulation run begins with all foragers starting at the nest (at the center of the world), and a newly generated resource distribution. After a simulation run starts, it continues for 2000 discrete timesteps. During each time step, all the agents (foragers, resources and patches) are updating and changing their state (if applicable). Foragers are either exploring for or exploiting the resources, communicating the resource locations or changing their activity due to activation-inactivation probabilities, errors and/or communication. Various output variables are calculated at the end of each simulation run, which are either an aggregate and/or averages of various other variables calculated during each run.

### Design concepts

#### Basic principles

The rationale for our modeling effort was that phenomena or behaviors at the level of individuals may have counterintuitive effects at the level of the collective, especially when communication or other interactions between individuals in a group allow for feedback loops [20,62–67]. It is thus a relevant and interesting question to explore which potential individual errors have the highest impact (whether or not real social insects frequently make these errors), and how already-known communication strategies in social insects may exacerbate or mitigate such impacts. With the insights from our model, we can better understand evolution of both individual accuracy (which errors are under selection pressure to avoid) and collective robustness (which communication types may evolve, in part, to mitigate errors).

*Emergence*

We expect that collective outcomes may not be intuitive even from known processes at the individual level. This phenomenon is generally termed ‘emergence’ [68,69]. In particular, all of the ‘performance measures’ used here are not built-in (See ‘Response variables quantified’ in the main manuscript), but are collective outcomes that are a result of repeated and recursive interactions between individuals and their environment (i.e. feedback loops). In addition to this classic view of ‘emergence’, theory in biology often serves to more precisely and quantitatively describe processes, the descriptions of which previously were verbal or intuited. Quantitative descriptions often highlight problems, inconsistencies, or surprising relationships or outcomes that are not apparent from apparently equivalent verbal descriptions [70,71].

*Adaptation*

No ‘adaptation’ or evolution is modeled here. Foragers do have ‘memory’, in that a single location of a resource is ‘learned’, either through direct experience (encountering a resource within the *sense-radius* while searching) or through communication (copied from another individual). We also do not model a dynamically changing environment, except in the sense that resources become exhausted over time. Any change in collective behavior in response to the initial distribution of resources or to the depletion of resources during the simulation is thus an emergent result of the interactions of individual foragers, directly or indirectly, their memory of resource locations, and their behavior rules (Fig. 1).

*Objectives*

As stated above, any 'objectives' that foragers have are only to the extent that they are implicit in the behavior represented in Fig. 1. No active maximization, evolution, or goal-seeking in foragers in addition to this behavior is modeled here. Foragers move in a correlated random walk when searching (the parameters of which are fixed). Whenever communication (not via pheromones) occurs, the information-receiving individual compares the communicated resource with the resource in its own memory, and retains a memory of the resource with the location closer to the nest. It can be said that individuals thus rate resources by a single quality dimension, distance to the (home) nest (at the center of the map). Implicitly, one might thus conclude that the individuals and the group aim to forage from the closest resources to the nest. If the communication type is via 'beacon' or 'trail' pheromones, foragers turn into the direction of detected pheromone; in 'trail' communication they do not differentiate between high or low levels of pheromone, and even in 'beacon' communication, no information about the quality of the resource is explicitly contained in the signal. Evaporation of pheromone over time may nonetheless lead to collective preference for closer resources [72]. We examine the implications of this implementation of pheromone recruitment further (See the 'Discussion' section in the main manuscript). In general, real food resources used by insects differ in other aspects of 'quality', and reacting to this may be a key benefit of communication [50,73–75], but this is not modeled here.

*Learning*

All foragers either start as explorers or inactive within the nest, in either case with no memory or (explicit) knowledge about the environment. As discussed under '*Adaptation*', foragers may have a memory of a single resource location, until they go to that location and find the resource empty. This location memory can also change if overwritten in the communication process (by an apparently ‘superior’ location which is closer to the nest), or if subject to errors. For instance, ‘wrong target memory’ error results in an incorrect location being remembered instead of refreshing the (accurate) current memory. Similarly, due to ‘forgetting errors’,at any time during a trip towards the location (i.e. while ‘exploiting') the memory may be ‘forgotten’. Other than this resource (coordinate) memory, there is no learning or memory present in the model.

*Prediction*

This aspect is not applicable for our model. The agents in our model are neither coded to (explicitly) predict future conditions nor consequences of their decisions.

*Sensing*

Foragers can sense the presence of a resource, other foragers, and the nest location only within their ‘*sense radius*’, and only do so when in a state that requires this (e.g. exploiters do not explore for, and thus sense, other resources on the way to a known resource location).

*Interaction*

Foragers in our model interact directly through communication, and indirectly through changes in the resource environments. Communication, in our model, is typically implemented as directly altering the receiving forager’s memory (typically by copying the remembered location from the sender). Foragers interact directly, through communication, and indirectly, through affecting the common environment (by depleting resources). In general, indirect interactions are likely to be important drivers of collective behavior [68,69,76]. Even though such indirect interactions are limited in scope in our model, biased exploitation of resources (resources near the nest are likely to be discovered and emptied faster) is likely to be an important process driving the overall distribution of foragers on the map as well as their efficiency. Additionally, the common environment is also affected by the deposition of pheromones when the trail or the beacon communication is present. In all other ways, actions by individual foragers are independent of actions of other foragers. We do not model physical collisions or crowding, because in our model, foragers are thought of much smaller in relation to the spatial map in which they move. We implemented five different communication 'architectures' or 'types', as well as 'no communication':

a. No communication: No direct interaction or information exchange between foragers.
b. Direct communication: Foragers (both explorers and exploiters) check on each time step with probability *signal-strength* whether at least one forager with a resource location memory is present in their *sense-radius*. If so, one of these foragers (potential senders of information) is chosen randomly, and their target memory is copied if this new location is nearer to the nest than the current location in the receiver's memory. A 'sender' may thus communicate with multiple receivers in the same time step, but a receiver will not be available to communicate with more than one sender per time step [77]. In ants, for example, this type of communication can occur when searching ants receive food from returning ants in the field via trophallaxis [78] or via tandem runs [79]. The received food may contain information about scent and quality of the resource, which is communicated in this food exchange and helps recruit the receivers to specific food sources [78].

c. Central-place communication: Foragers in the nest (inactive foragers) extract the resource location from other foragers within their *sense-radius*. This is thus similar to 'direct' communication, but only taking place in the nest itself. For example, honey bees (*Apis sp.*) foragers communicate distance and direction of food sources using the 'waggle dance' on comb in their nest [80].

d. Blackboard communication: The nest extracts the nearest remembered resource location among all other remembered resources, and sets it as the nest-memory. Inactive foragers at the nest probabilistically copy this nest memory. In bumble bees (*Bombus terrestris*), for example, the honey storage within the nest acts as an information repository - more bees begin foraging when the honey pot levels begin to fill, and if the nectar quality levels deposited by the incoming workers are high [81]. Furthermore, the foragers can learn specific floral odors when encountering incoming nestmates, and visit those specific floral patches for the nectar [82,83].

e. Trail communication: All foragers returning to the nest with food probabilistically deposit pheromone. These pheromone deposits can thus form a kind of 'breadcrumb trail' from the nest to the food. This trail is disappearing at a constant rate (empirically, because the volatile pheromone evaporates; in our model, each time step the amount of pheromone on each patch is reduced to *pheromone-deposited x* (1- *pheromone-fade-rate*).

Both in *beacon* and *trail* communication, the walking direction (i.e. turning angles) of explorers are affected by pheromone levels in a cone determined by the *tortuosity* and *sense-radius* parameters both to the right and left side. If no pheromone is detected, a normal random walk is used. If the pheromone concentration is high enough on both sides, they walk straight. If pheromone is detectable on one but not the other side, they turn to that side at an angle of *tortuosity*/2. This is similar to how ants use their antennae to detect the trail, and use the differences in the concentration detected by both antennae to turn towards that direction (tropotaxis) [38,84,85], but it is important to note that in our model the ants do not actually compare the pheromone concentrations quantitatively (other than to detect presence or absence of detectable pheromone levels). Explorers thus following a trail or pheromone field do not know at what coordinate to expect a food source, and thus are available to detect any food source once they are within sense-radius. Once a food source is detected, these explorers save the coordinate in memory and become exploiters. Once a forager has detected a food source, it uses its memory to return (and the pheromone is no longer relevant). In reality, ant species vary in their reliance on the pheromone trail versus their own memory, or on a combination of both information sources [48,86,87].

f. Beacon communication: A forager (an exploiter) that collects food from a resource may deposit pheromone at the food patch before returning to the nest. The pheromone serves to attract other nearby explorers to the resource. For example, stingless bees (e.g. *Melipona panamica*) deposits a volatile pheromone on food sources that attracts nearby foragers [88]. In our model, once a patch contains any pheromone, the pheromone diffuses into the eight neighboring patches on each time step (with each neighbor receiving 1/8th of the total pheromone concentration). At the same time, the pheromone level in each patch reduces at a constant rate (*pheromone-fade-rate*) each time step, and as a result it never exhausts completely. However, if the pheromone concentration in the cone (determined by *tortuosity* and *sense-radius*, see above in the trail communication) is less than 50 *pheromone-deposited* units on either side, then it becomes too low to be detected.

*Stochasticity*

Our model is not deterministic, and many actions and outcomes of actions are probabilistic. As a result, we see a high degree of variance in outcome variables even for constant input parameter values. A major source of variation in outcomes is that the exact positions of resources are randomized for every simulation run. For example, the distance of the closest resource patch may vary substantially between simulations (see panel d in Fig. 7, and Figs. S8-12. Explorers also move in a (correlated) random walk in their environment to search for resources. Many state transitions are probabilistic, such as inactive foragers becoming active with an activation probability.

*Collectives*

The foragers together form a collective (i.e., a colony). Our model assumes that all resources are collected to a single point (the nest). All of the outcome variables we measure are group-level outputs. Our model is thus most appropriately seen as representing a social insect colony that is interested in maximizing colony-level food collection [68,89].

*Observation*

All outcomes of simulations analyzed were only quantified once at the end of each simulation run, although some of them required the ‘world observer’ in Netlogo to keep track of accumulating information (such as the number of trips performed) throughout the run. We quantified four categories of performance measures- (1) Overall foraging performance, (2) exploration, (3) exploitation, and (4) relative performances. For overall performance measure, we quantified total resources collected by the whole colony during each simulation run (e.g., total food (seeds or nectar) collected during a whole day of foraging -[67,90]. We quantified exploration to quantify the time investment and search efficiency of foragers. Specifically, we quantified resources discovered by the whole colony (how many resources the colony discovered during the whole foraging day e.g., [91]), total forager-time spent searching (Total time invested by foragers in searching for resources; e.g., [92,93]), the area explored per unit time spent by foragers in searching for resources (The extent to which the explored area overlapped between foragers; e.g., [94,95], and search efficiency (resources discovered per unit time spent by foragers e.g., [91]. For quantifying exploitation, we focussed on resource units collected per resource patch discovered (return on energetic investment during foraging e.g., [96], resources collected per unit time spent collecting resources (resource collection efficiency; e.g., [97], and time spent collecting resources per unit resource discovered (Return on time investment for every resource unit discovered, e.g., [97]).

We also quantify the impact of different failure rates (error probability). Such robustness analyses are done in material science to examine the behavior of a material under increasing stress [98]. Moreover, from the characteristic shape of the curve, one can infer if the material is ductile or brittle against the given stress [98]. Specifically, for each combination of error and communication type, we quantified relative performance by comparing resource units collected relative to median resource units collected without errors (i.e., 100% functionality). This allowed us to standardize the error-free performance as 1, and measure the proportional decrease in performance under increasing probability of errors.

Additionally, we recorded a number of other properties of simulation runs and colony behavior, as shown in Fig. 6. Specifically, we quantified how resource collection varied due to a variety of variables such as proportion of time spent by foragers in different activities (searching and collecting), time at which which they discovered the first resource, total area explored, average distance of the resource from the nest, total trips taken by foragers (search and collection combined), efficiency, number of resources discovered, average search distance of foragers, average distance of an exploited resource and number of unsuccessful search trips. All these measures were quantified at the colony-level to further uncover the effects of errors on exploration and exploitation across different communication types.

8. Exélis MP, Ramli R, Abdul Latif SA, Idris AH, Clemente-Orta G, Kermorvant C. Elucidating the daily foraging activity pattern of Oecophylla smaragdina to minimize bite

nuisances in Asia large agro-system plantations. Heliyon. 2024;10: e26105. doi:10.1016/j.heliyon.2024.e26105

48. Lanan M. Spatiotemporal resource distribution and foraging strategies of ants (Hymenoptera: Formicidae). 2014.

80. Frisch K von. Dance language and orientation of bees. 1967.

81. Dornhaus A, Chittka L. Bumble bees (Bombus terrestris) store both food and information in honeypots. Behav Ecol. 2005;16: 661–666. doi:10.1093/beheco/ari040

82. Dornhaus A, Chittka L. Evolutionary origins of bee dances. Nature. 1999;401: 38–38. doi:10.1038/43372

83. Molet M, Chittka L, Raine NE. How floral odours are learned inside the bumblebee (Bombus terrestris) nest. Naturwissenschaften. 2009;96: 213–219. doi:10.1007/s00114-008-0465-x

84. Hölldobler B, Wilson EO. The ants. Harvard University Press; 1990.
