## Supplementary materials (R code, model description and additional plots) for "Frequent errors are the worst: robustness to individual failures in collective foraging": Supplementary file S2_2026.pdf

### Supplementary figures & tables

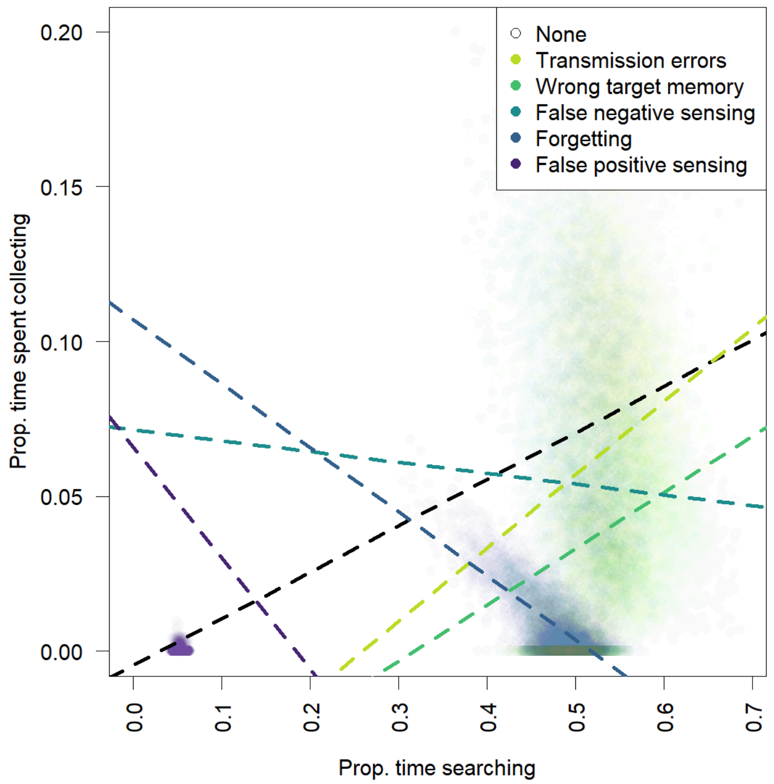


Fig. S1: Illustrated here just for central-place communication, the relationship between time invested in searching and time then spent collecting from discovered resources generally did not trade off, instead more time spent searching led to more resources discovered and thus more time spent collecting. The exception to this were the cases with more severe errors: in false positive sensing, a lot of time was spent 'collecting' but without actually arriving to the (data points between 0.2 and 0.6 not shown in this graph); with forgetting, there is a clear negative relationship shown here, with the variation probably explained by differences in how close the closest resources happened to be in the environment. For false negative sensing, the estimated relationship is also slightly negative (although individual simulations vary widely).

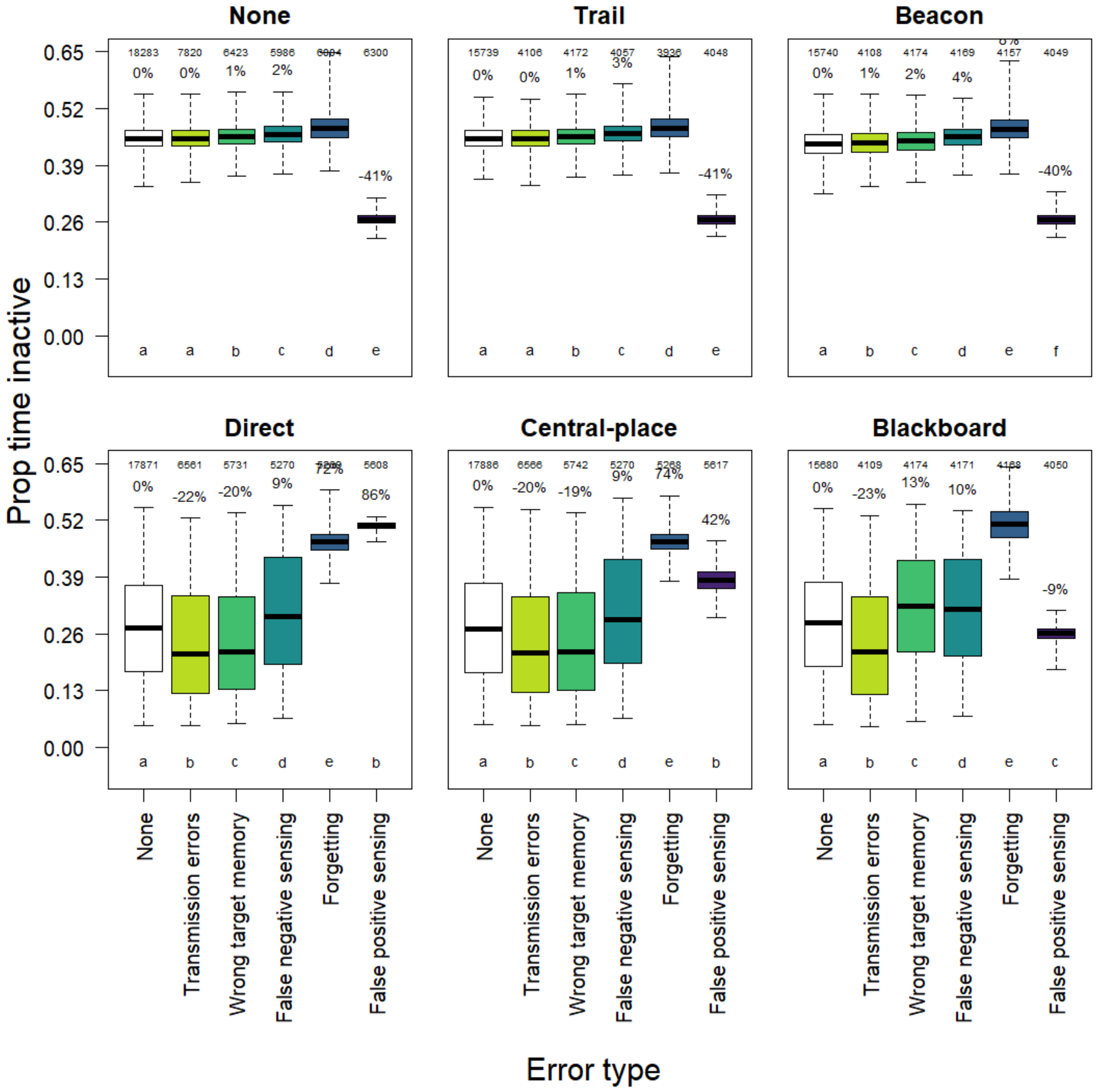


Fig S2: Proportion of time spent inactive by the colony plotted against error types separately for each communication type. All the box plots here show the median (dark black line), quartiles, and the range of the total resources collected for each error type. The numbers at the top of the graph indicate the sample size (number of simulation runs) for each error type. The bars that do not share the same letter (a-e) are significantly different from each other (significant Kruskal-Wallis test (p<0.05) followed by Dunn's post hoc test, p<0.05). The percentages above the bars within a panel indicate median performance relative to the median value without errors (white bar). Forgetting errors leads to the highest increase in the proportion of time spent inactive.

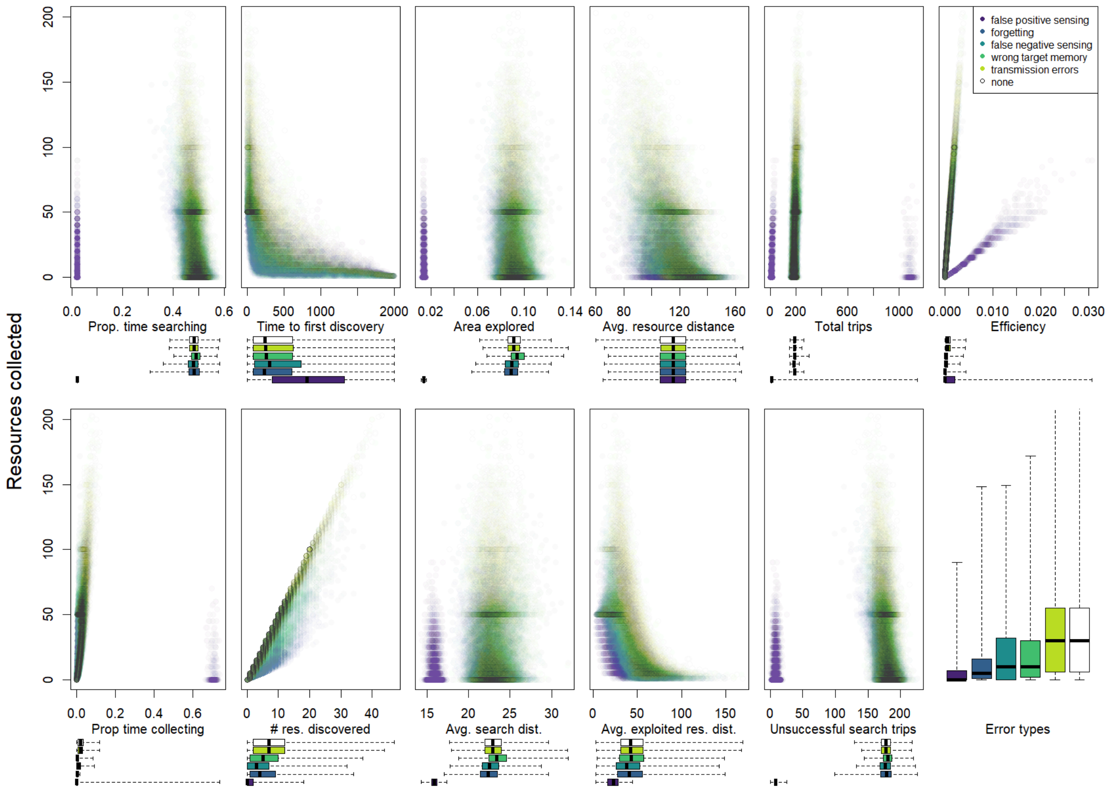


Fig. S3: Impact of errors on different measures of colony performance and behavior for no communication. Boxplots underneath scatterplots show x-axis value distributions for each error type; y-axis (performance) distributions for each error type are shown in panel l (and are identical across all scatterplots; panel l matches corresponding data in Fig. 2). Dark horizontal lines show high point densities; since resources occurred in 'clusters' of 10 resource patches carrying 5 resource units each, colonies that emptied exactly one resource cluster collected 50 resource units (or multiples of 50 for multiple clusters). Intermediate values indicate simulation runs in which some resource clusters were not fully exploited.

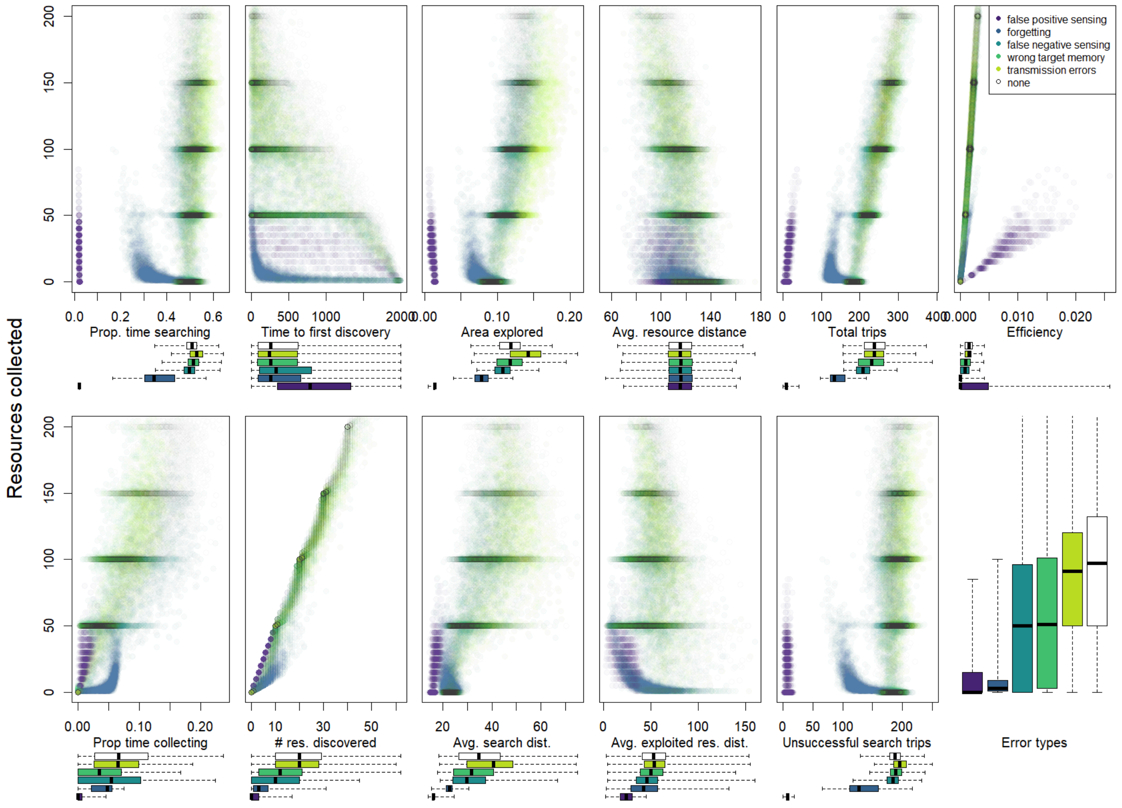


Fig. S4: Impact of errors on different measures of colony performance and behavior for blackboard communication. Boxplots underneath scatterplots show x-axis value distributions for each error type; y-axis (performance) distributions for each error type are shown in panel I (and are identical across all scatterplots; panel I matches corresponding data in Fig. 2 (top six panels)). Dark horizontal lines show high point densities; since resources occurred in ‘clusters’ of 10 resource patches carrying 5 resource units each, colonies that emptied exactly one resource cluster collected 50 resource units (or multiples of 50 for multiple clusters). Intermediate values indicate simulation runs in which some resource clusters were not fully exploited.

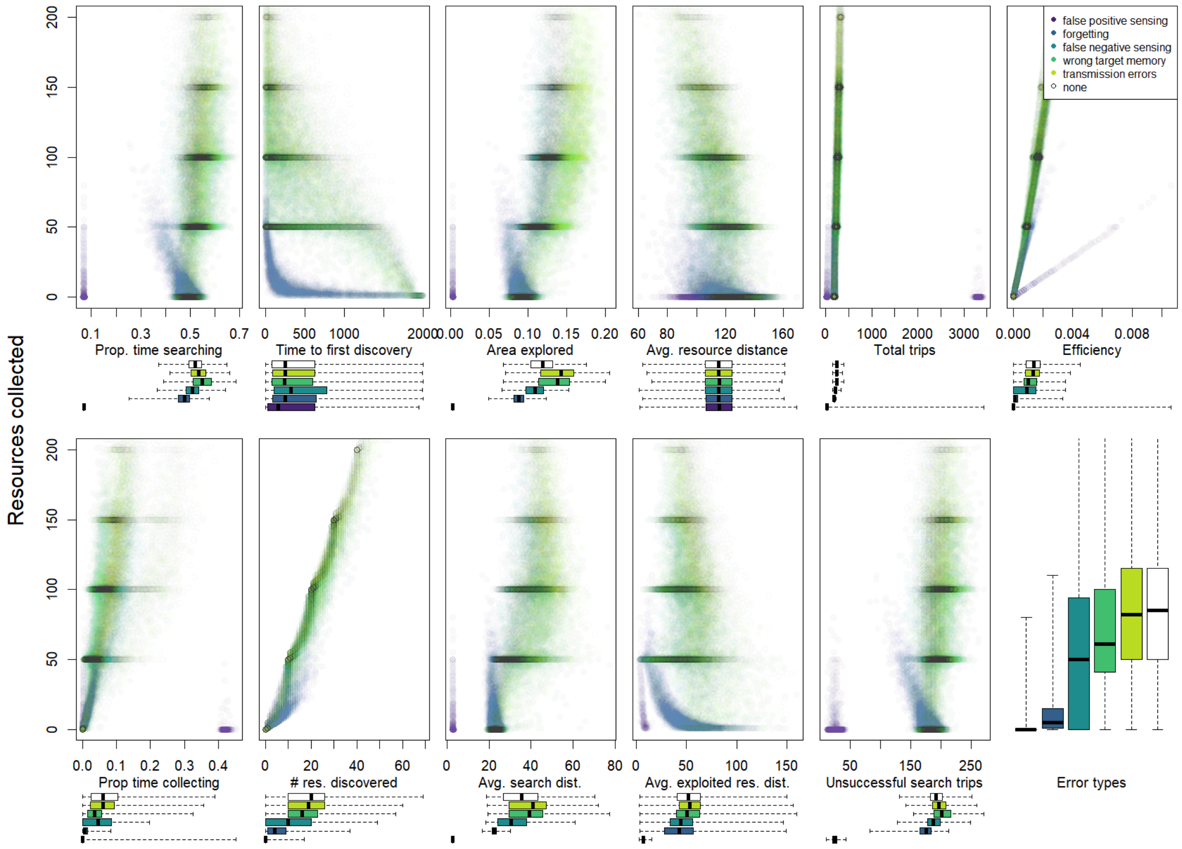


Fig. S5: Impact of errors on different measures of colony performance and behavior for direct communication. Boxplots underneath scatterplots show x-axis value distributions for each error type; y-axis (performance) distributions for each error type are shown in panel I (and are identical across all scatterplots; panel I matches corresponding data in Fig. 2 (top six panels)). Dark horizontal lines show high point densities; since resources occurred in 'clusters' of 10 resource patches carrying 5 resource units each, colonies that emptied exactly one resource cluster collected 50 resource units (or multiples of 50 for multiple clusters). Intermediate values indicate simulation runs in which some resource clusters were not fully exploited.

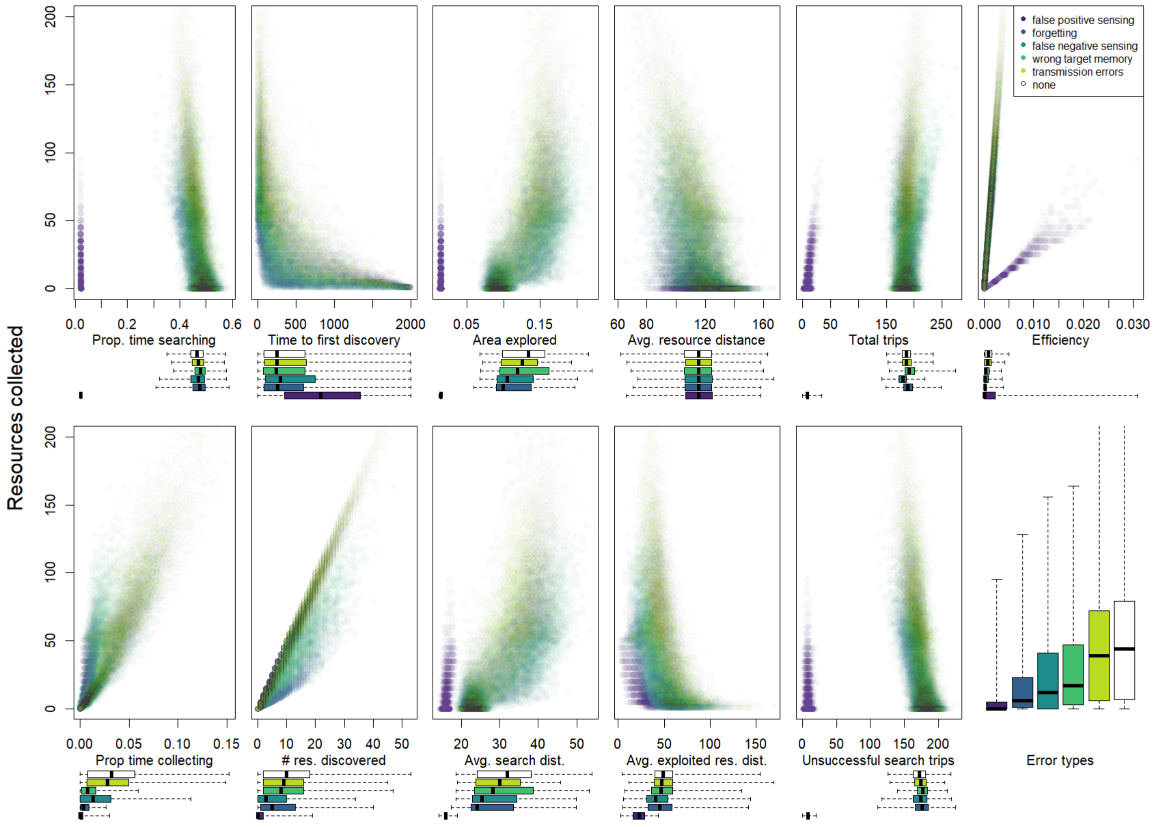


Fig. S6: Impact of errors on different measures of colony performance and behavior for beacon communication. Boxplots underneath scatterplots show x-axis value distributions for each error type; y-axis (performance) distributions for each error type are shown in panel I (and are identical across all scatterplots; panel I matches corresponding data in Fig. 2 (top six panels)). Dark horizontal lines show high point densities; since resources occurred in 'clusters' of 10 resource patches carrying 5 resource units each, colonies that emptied exactly one resource cluster collected 50 resource units (or multiples of 50 for multiple clusters). Intermediate values indicate simulation runs in which some resource clusters were not fully exploited.

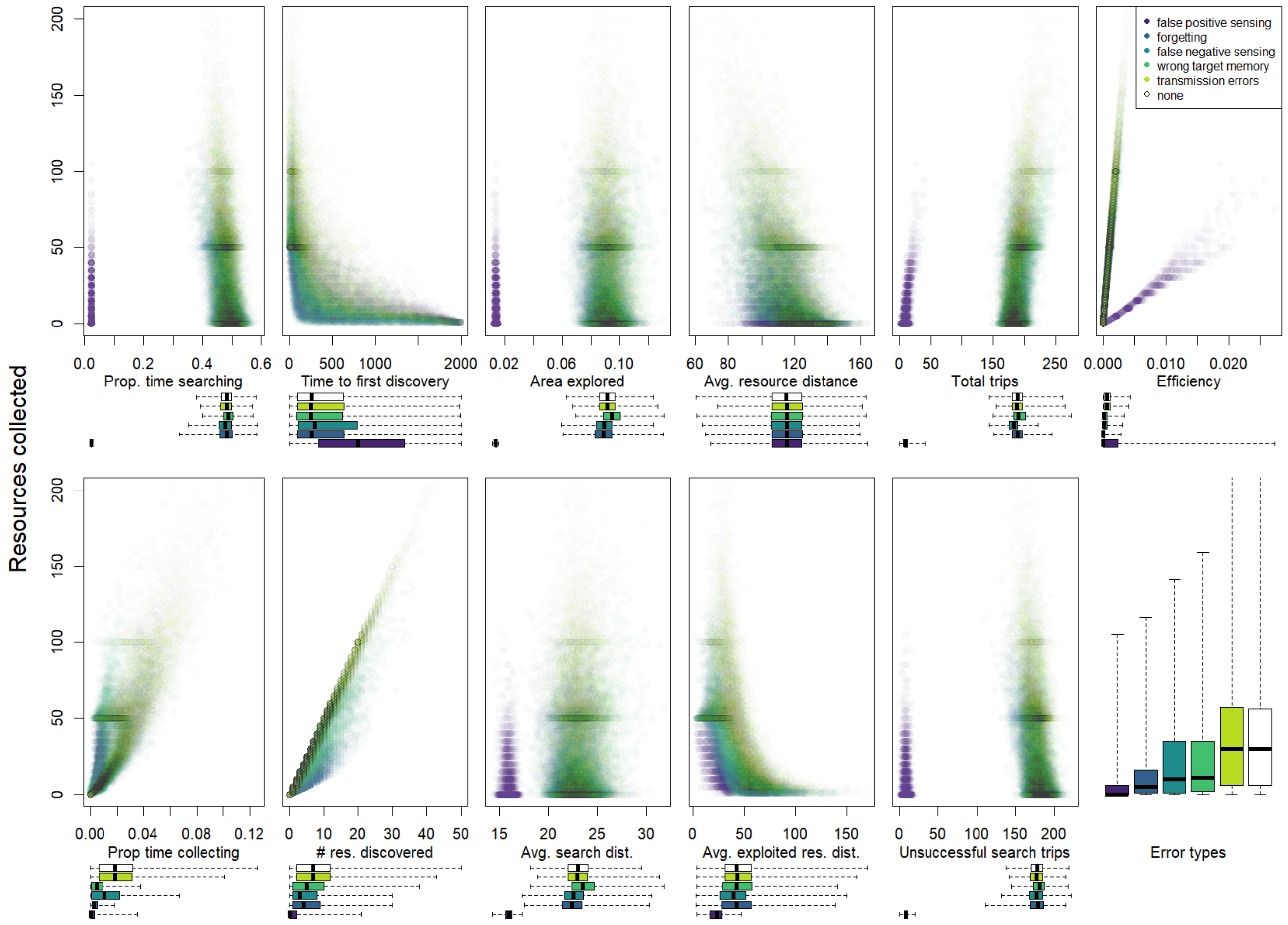


Fig. S7: Impact of errors on different measures of colony performance and behavior for trail communication. Boxplots underneath scatterplots show x-axis value distributions for each error type; y-axis (performance) distributions for each error type are shown in panel I (and are identical across all scatterplots; panel I matches corresponding data in Fig. 2 (top six panels)). Dark horizontal lines show high point densities; since resources occurred in 'clusters' of 10 resource patches carrying 5 resource units each, colonies that emptied exactly one resource cluster collected 50 resource units (or multiples of 50 for multiple clusters). Intermediate values indicate simulation runs in which some resource clusters were not fully exploited.

**Kruskal-Wallis test result tables**

Table S2: Kruskal Wallis test results for overall foraging performance (total resources collected) ~ error types for each communication type. Each Kruskal-Wallis test here had the same degrees of freedom (df = 5) and was statistically significant ($p < 0.001$). Dunn's post hoc test results are indicated by letters in Fig. 2.

| **Communication type** | **Sample size** | $\chi^2$ | **p-value** |
|---|---|---|---|
| No communication | 50816 | 8863.4 | <0.001 |
| Trail communication | 36058 | 5983.7 | <0.001 |
| Beacon communication | 36397 | 6636.6 | <0.001 |
| Direct communication | 46310 | 15127.9 | <0.001 |
| Central place communication | 46349 | 14810 | <0.001 |
| Blackboard communication | 36352 | 10146.4 | <0.001 |

Table S3: Kruskal Wallis test results for overall foraging rate (total resources collected) ~ communication type for each error type. Each Kruskal-Wallis test here had the same degrees of freedom (df = 5) and was statistically significant ($p < 0.001$). Dunn's post hoc test results are indicated by letters in Fig. X.

| **Communication type** | **Sample size** | $\chi^2$ | **p-value** |
|---|---|---|---|
| No error | 101199 | 13598 | <0.001 |
| Transmission errors | 33270 | 4658.7 | <0.001 |
| Wrong target memory | 30416 | 5573 | <0.001 |
| False negative sensing | 28923 | 3445.5 | <0.001 |
| Forgetting errors | 28802 | 235.8 | <0.001 |
| False positive sensing | 29672 | 2791.5 | <0.001 |

Table S4: Kruskal Wallis test results for total resources discovered ~ error types for each communication type. Each Kruskal-Wallis test here had the same degrees of freedom (df = 5) and was statistically significant ($p < 0.001$). Dunn's post hoc test results are indicated by letters in Fig. 4.

| **Communication type** | **Sample size** | $\chi^2$ | **p-value** |
|---|---|---|---|
| No communication | 50816 | 7642.4 | <0.001 |
| Trail communication | 36058 | 5134.9 | <0.001 |
| Beacon communication | 36397 | 5842.6 | <0.001 |
| Direct communication | 46310 | 14064 | <0.001 |
| Central place communication | 46349 | 13719 | <0.001 |
| Blackboard communication | 36352 | 9685.7 | <0.001 |

Table S5: Kruskal Wallis test results for total resources discovered per unit area explored (search efficiency) ~ error types for each communication type. Each Kruskal-Wallis test here had the same degrees of freedom (df = 5) and was statistically significant ($p < 0.001$). Dunn's post hoc test results are indicated by letters in Fig. 4.

| **Communication type** | **Sample size** | $\chi^2$ | **p-value** |
|---|---|---|---|
| No communication | 50816 | 2737.4 | <0.001 |
| Trail communication | 36058 | 1898.5 | <0.001 |
| Beacon communication | 36397 | 2051.5 | <0.001 |
| Direct communication | 46310 | 10506.1 | <0.001 |
| Central place communication | 46349 | 9680.8 | <0.001 |
| Blackboard communication | 36352 | 3283.5 | <0.001 |

Table S6: Kruskal Wallis test results for proportion of patches explored per unit time spent searching (Avoidance of area overlap between foragers) ~ error types for each communication type. Each Kruskal-Wallis test here had the same degrees of freedom (df = 5) and was statistically significant ($p < 0.001$). Dunn's post hoc test results are indicated by letters in Fig. 4.

| **Communication type** | **Sample size** | $\chi^2$ | **p-value** |
|---|---|---|---|
| No communication | 50816 | 17416.2 | <0.001 |
| Trail communication | 36058 | 11437.3 | <0.001 |
| Beacon communication | 36397 | 11757.6 | <0.001 |
| Direct communication | 46310 | 22704.9 | <0.001 |
| Central place communication | 46349 | 22756.9 | <0.001 |
| Blackboard communication | 36352 | 12738.6 | <0.001 |

Table S7: Kruskal Wallis test results for the proportion of time spent searching by foragers ~ error types for each communication type. Each Kruskal-Wallis test here had the same degrees of freedom (df = 5) and was statistically significant ($p < 0.001$). Dunn's post hoc test results are indicated by letters in Fig. 4.

| **Communication type** | **Sample size** | $\chi^2$ | **p-value** |
|---|---|---|---|
| No communication | 50816 | 17002.3 | <0.001 |
| Trail communication | 36058 | 11059.3 | <0.001 |
| Beacon communication | 36397 | 11240.9 | <0.001 |
| Direct communication | 46310 | 21260.3 | <0.001 |
| Central place communication | 46349 | 21011.7 | <0.001 |
| Blackboard communication | 36352 | 18081 | <0.001 |

Table S8: Kruskal Wallis test results for the proportion of time spent by foragers collecting resources per unit resource discovered (Investment in exploitation)~ error types for each communication type. Each Kruskal-Wallis test here had the same degrees of freedom (df = 5) and was statistically significant ($p < 0.001$). Dunn's post hoc test results are indicated by letters in Fig. 5.

| **Communication type** | **Sample size** | $\chi^2$ | **p-value** |
|---|---|---|---|
| No communication | 39041 | 20628.9 | <0.001 |
| Trail communication | 27929 | 14624.9 | <0.001 |
| Beacon communication | 28079 | 16373.6 | <0.001 |
| Direct communication | 34209 | 10767.8 | <0.001 |
| Central place communication | 34195 | 12822.6 | <0.001 |
| Blackboard communication | 28049 | 10729.7 | <0.001 |

Table S9: Kruskal Wallis test results for the total collected resources per unit resource discovered (Exploitation efficiency) ~ error types for each communication type. Each Kruskal-Wallis test here had the same degrees of freedom (df = 5) and was statistically significant ($p < 0.001$). Dunn's post hoc test results are indicated by letters in Fig. 5.

| **Communication type** | **Sample size** | $\chi^2$ | **p-value** |
|---|---|---|---|
| No communication | 39041 | 17400.9 | <0.001 |
| Trail communication | 27929 | 11493.5 | <0.001 |
| Beacon communication | 28079 | 13079.1 | <0.001 |
| Direct communication | 34209 | 13195.6 | <0.001 |
| Central place communication | 34195 | 13362.3 | <0.001 |
| Blackboard communication | 28049 | 10874.5 | <0.001 |

Table S10: Kruskal Wallis test results for the total collected resources per unit time spent collecting (Collection efficiency) ~ error types for each communication type. Each Kruskal-Wallis test here had the same degrees of freedom (df = 5) and was statistically significant ($p < 0.001$). Dunn's post hoc test results are indicated by letters in Fig. 5.

| **Communication type** | **Sample size** | $\chi^2$ | **p-value** |
|---|---|---|---|
| No communication | 38854 | 8379.9 | <0.001 |
| Trail communication | 27563 | 6592.6 | <0.001 |
| Beacon communication | 27806 | 10960.2 | <0.001 |
| Direct communication | 34515 | 5254.1 | <0.001 |
| Central place communication | 34498 | 3785.3 | <0.001 |
| Blackboard communication | 27696 | 11231.1 | <0.001 |

Table S11: Kruskal Wallis test results for the proportion of time spent inactive by foragers ~ error types for each communication type. Each Kruskal-Wallis test here had the same degrees of freedom (df = 5) and was statistically significant ($p < 0.001$). Dunn's post hoc test results are indicated by letters in Fig. S2.

| **Communication type** | **Sample size** | $\chi^2$ | **p-value** |
|---|---|---|---|
| No communication | 50816 | 19044.8 | <0.001 |
| Trail communication | 36058 | 12692.3 | <0.001 |
| Beacon communication | 36397 | 14063.1 | <0.001 |
| Direct communication | 46310 | 22157.5 | <0.001 |
| Central place communication | 46349 | 13753.2 | <0.001 |
| Blackboard communication | 36352 | 10806.2 | <0.001 |
